# Minimal effects of proto-Y chromosomes on house fly gene expression in spite of evidence that selection maintains stable polygenic sex determination

**DOI:** 10.1101/545178

**Authors:** Jae Hak Son, Tea Kohlbrenner, Svenia Heinze, Leo Beukeboom, Daniel Bopp, Richard P. Meisel

## Abstract

Sex determination, the developmental process by which organismal sex is established, evolves fast, often due to changes in the master regulators at the top of the pathway. Additionally, in species with polygenic sex determination, multiple different master regulators segregate as polymorphisms. Understanding the forces that maintain polygenic sex determination can be informative of the factors that drive the evolution of sex determination. The house fly, *Musca domestica*, is a well-suited model to those ends because natural populations harbor male-determining loci on each of the six chromosomes and a bi-allelic female-determiner. To investigate how natural selection maintains polygenic sex determination in house fly, we assayed the phenotypic effects of proto-Y chromosomes by performing RNA-seq experiments to measure gene expression in house fly males carrying different proto-Y chromosomes. We find that the proto-Y chromosomes have similar effects as a non-sex-determining autosome. In addition, we created sex-reversed males without any proto-Y chromosomes, and they had nearly identical gene expression profiles as genotypic males. Therefore, the proto-Y chromosomes have a minor effect on male gene expression, consistent with previously described minimal X-Y sequence differences. Despite these minimal differences, we find evidence for a disproportionate effect of one proto-Y chromosome on male-biased expression, which could be partially responsible for fitness differences between males with different proto-Y chromosome genotypes. Our results therefore suggest that, if natural selection maintains polygenic sex determination in house fly via gene expression differences, the phenotypes under selection likely depend on a small number of genetic targets.

## Introduction

Sex determination is the process by which genetic or environmental cues cause an individual to develop into either a female or male. Sex determination evolves rapidly, often due to changes in the master regulatory genes at the top of sex determining pathways (Beukeboom and Perrin 2014). Sex determining pathways can also be variable (polygenic or multifactorial) within species (Moore and Roberts 2013). Many population genetic models predict that polygenic sex determination should be a transient state between monogenic equilibria (Rice 1986; van Doorn and Kirkpatrick 2007). It is therefore surprising that polygenic sex determination has been found in numerous species (Orzack *et al*. 1980; Moore and Roberts 2013; Bachtrog *et al*. 2014). Understanding how polygenic sex determining systems are maintained will help shed light on the forces driving the rapid evolution in sex determination pathways.

The house fly, *Musca domestica*, is a model species to study polygenic sex determination because it has a well characterized and highly variable sex determination system (Dübendorfer *et al*. 2002; Hamm *et al*. 2015). The male-determining gene, *Mdmd*, appears to be recently derived in house fly as it is absent in its close relative *Stomoxys calcitrans*, and it cannot be found in other related dipterans (Sharma *et al*. 2017). *Mdmd* regulates the splicing of the house fly ortholog of *transformer* (*Md-tra*), preventing males from producing a functional female-determining isoform of *Md-tra* (Hediger *et al*. 2010; Sharma *et al*. 2017). A dominant female-determining allele (*Md-tra^D^*) that is not sensitive to *Mdmd* regulation also segregates in natural populations, allowing females to carry *Mdmd* (McDonald *et al*. 1978; Kozielska *et al*. 2008; Hediger *et al*. 2010).

*Mdmd* can be found on multiple different chromosomes in natural populations of house fly (Sharma *et al*. 2017). The two most common locations of *Mdmd* in natural populations are the Y chromosome (Y^M^) and third chromosome (III^M^) (Hamm *et al*. 2015). Historically, the chromosomes carrying the male determiner were designated as the Y (Y^M^), X (X^M^), and any of the five autosomes (e.g., III^M^). However, recent work showed that the Y^M^ chromosome is highly similar in gene content to the X chromosome because Y^M^ is a very young proto-Y chromosome (Meisel *et al*. 2017). These findings align with the independent discovery that *Mdmd* is of a recent origin (Sharma *et al*. 2017). Moreover, previous work observed minimal morphological and sequence differences between the X and Y chromosomes (Boyes *et al*. 1964; Hediger *et al*. 1998). The third chromosome carrying *Mdmd* is also very recently derived from the standard third chromosome (Meisel *et al*. 2017). We therefore refer to any chromosome carrying *Mdmd* (including Y^M^ and III^M^) as a proto-Y chromosome.

There are multiple lines of evidence that natural selection maintains polygenic sex determination in house fly. First, Y^M^ and III^M^ form stable latitudinal clines on multiple continents (Franco *et al*. 1982; Tomita and Wada 1989; Hamm *et al*. 2005; Kozielska *et al*. 2008). Y^M^ is most frequent in northern populations, and III^M^ predominates in southern populations. The distributions of the proto-Y chromosomes correlate with seasonality in temperature (Feldmeyer *et al*. 2008), suggesting that temperature modulates the fitness of males carrying different proto-Y chromosomes. Second, males carrying Y^M^ or III^M^ differ in their success courting female mates, and the frequency of the III^M^ chromosome reproducibly increased over generations in laboratory population cages kept at a warm temperature (Hamm *et al*. 2009). Third, in some populations, individual males carry multiple proto-Y chromosomes, which would cause them to produce male-biased broods with their mates (Kozielska *et al*. 2006; Hamm *et al*. 2015). The frequency of males that carry multiple proto-Y chromosomes is positively correlated with the frequency of *Md-tra^D^* across populations (Meisel *et al*. 2016). This suggests that *Md-tra^D^* invaded to balance the sex ratio or *Md-tra^D^* allows for the increase in frequency of proto-Y chromosomes.

If natural selection maintains polygenic sex determination in house fly, Y^M^ and III^M^ must have different phenotypic and fitness effects for selection to act upon. However, a recent analysis of Y^M^ and III^M^ sequences revealed very few differences from their homologous X and III chromosomes, respectively (Meisel *et al*. 2017). To examine this paradox of evolutionarily important phenotypic effects of proto-Y chromosomes yet minimal sequence divergence from their homologs, we used high throughput mRNA sequencing (RNA-seq) to measure gene expression in house flies with different Y^M^ and III^M^ genotypes. This included testing the effects of multiple different naturally occurring Y^M^ and III^M^ chromosomes on a common genetic background. We also used RNA interference (RNAi) to knock down *Md-tra* and create sex-reversed males that do not carry any proto-Y chromosomes (Hediger *et al*. 2010), which we compared to males with the same genetic background carrying III^M^. Our experiments therefore allow us to identify candidate phenotypic differences between males carrying different proto-Y chromosomes upon which natural selection can act to maintain polygenic sex determination in house fly.

## Materials and Methods

### Strains with naturally occurring proto-Y chromosomes

We examined gene expression in four house fly strains that each have a different naturally occurring proto-Y chromosome (either Y^M^ or III^M^) on a common genetic background (Figure 1). We used a previously described backcrossing method to move each proto-Y chromosome onto the Cornell Susceptible (CS) genetic background (Meisel *et al*. 2015). CS is an inbred III^M^ strain produced by mixing strains collected from throughout the United States (Scott *et al*. 1996). Our first proto-Y chromosome is the III^M^ chromosome from the CS strain on its native background. The second strain (CSrab) was created by backcrossing the III^M^ chromosome from the rspin strain collected in New York (Shono and Scott 2003) onto the CS background for 10 generations, replacing the CS III^M^ chromosome (Supplementary Figure 1). The third strain (IsoCS) is a Y^M^ strain that was previously created by introducing the Y^M^ chromosome from a strain collected in Maine onto the CS background without III^M^ (Hamm *et al*. 2009). The fourth strain was created to test the effect of a non-*Mdmd*-bearing third chromosome on gene expression (Supplementary Figure 2). To that end, we introduced the third chromosome carrying the recessive *brown body* mutation (*bwb*) and the Y^M^ chromosome from the genome reference strain (aabys) onto the CS background to create the bwbCS strain (III*^bwb^*/ III*^bwb^*; X/Y^M^). We then crossed bwbCS males with CS females (bwbCS×CS) to create males that carry the aabys Y^M^ chromosome and are heterozygous for the non-*Mdmd* third chromosomes from CS and aabys on a CS background (III^CS^/ III*^bwb^*; X^CS^/Y^M^). We therefore have two III^M^ strains (CS and CSrab) with different origins of the III^M^ chromosome and two Y^M^ strains (IsoCS and bwbCS×CS) with different origins of the Y chromosome. In three of the strains (CS, CSrab, and IsoCS), females are isogenic for the CS background and males are isogenic except for their *Mdmd*-bearing proto-Y chromosomes.

**Figure 1.**
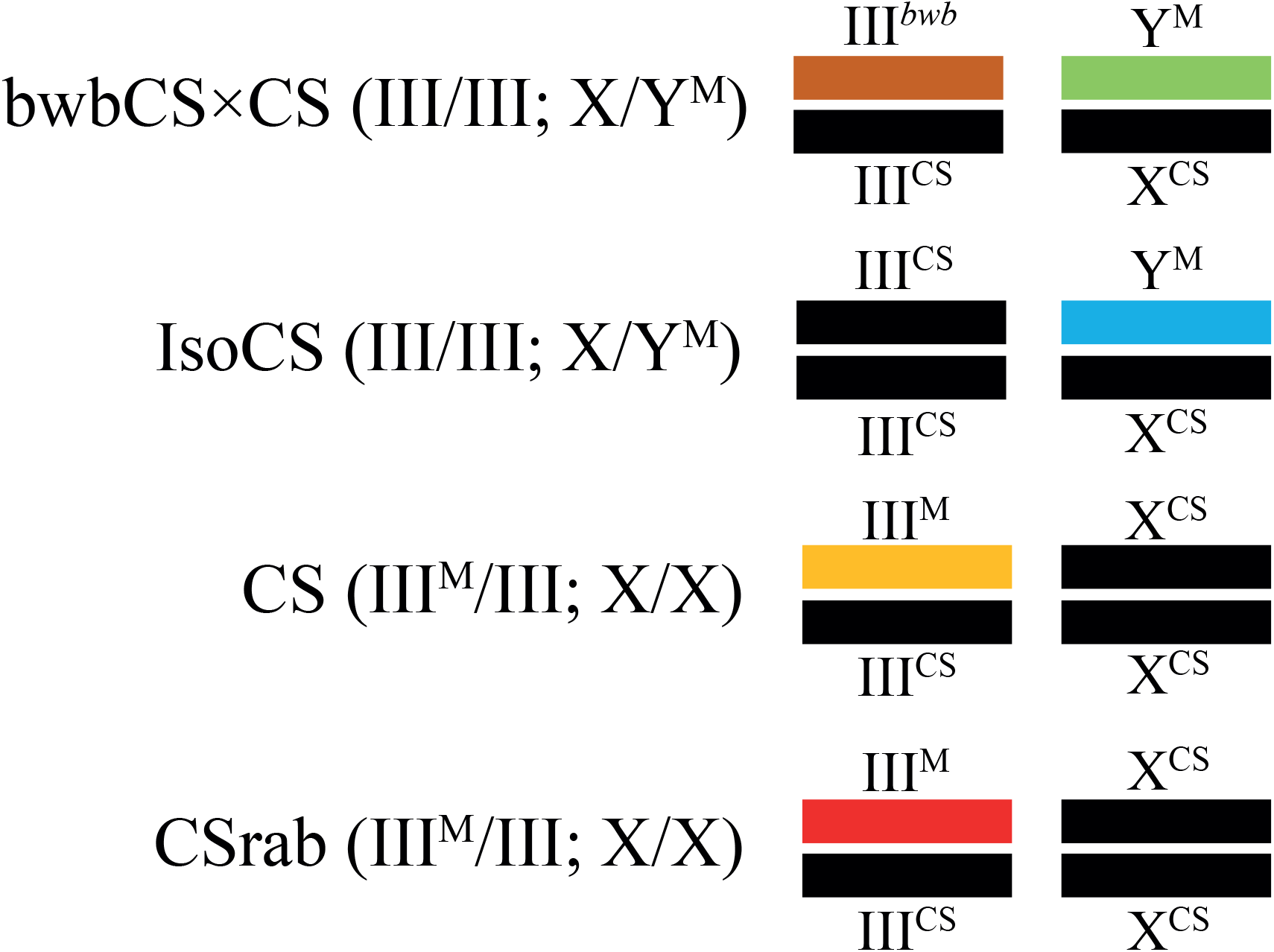
The genotypes of four strains with different naturally occurring Y^M^ or III^M^ proto-Y chromosomes on a common genetic background are shown. Black bars represent chromosomes from the common genetic background, and colored bars are chromosomes that are replaced on that background. Different colors of chromosomes indicate their origins from different strains. Chromosomes in the rest of genome (not shown), are from the common genetic background as well.

### RNAi knockdown to create sex reversed flies

We used RNAi targeting *Md-tra* to create sex-reversed males that do not carry a male-determining proto-Y chromosome. For the RNAi experiments we used a house fly strain that allows easy identification of sex-reversed individuals that are genotypic females but phenotypic males (Hediger *et al*. 2010). Females of this strain are homozygous for a third chromosome containing recessive alleles for *pointed wing* (*pw*) and *bwb*. Males carry one copy of the third chromosome with *pw* and *bwb*, and one copy of a III^M^ chromosome with wild-type alleles (*Mdmd pw*^+^ *bwb*^+^/*pw bwb*). Females therefore have pointed wings and brown bodies, as do sex-reversed males, whereas normal males have wild-type wings and wild-type bodies. We cannot do this same experiment with Y^M^ males because there are no morphological markers on the X/Y chromosome.

Double-stranded RNA (dsRNA) targeting *Md-tra* (*Md-tra-*RNAi) and GFP (GFP-RNAi) was generated and injected into early blastoderm embryos of the *pw bwb* strain following established protocols (Hediger *et al*. 2001, 2004). The fragment of dsRNA targeting *Md-tra* ranges from exon 1 to exon 5, and it was generated by amplifying cDNA from female house flies (Hediger *et al*. 2010). The sequences of the T7 extended primers used to produce dsRNA targeting *Md-tra* are 5’-gtaatacgatcactatagggTGGTGTAATATGGCTCTATCG-3’ and 5’-gtaatacgatcactatagggGCTGCCATACAAACGTGTC-3’ (sequences in lower case are the T7 region and sequences in upper case anneal to *Md-tra*). The sequences of the T7 extended primers used to produce dsRNA targeting GFP are 5’-gtaatacgatcactatagggATGTGAGCAAGGGC-3’ and 5’-gtaatacgatcactatagggCTTGTACAGCTCGTC-3’.

The larvae that hatched from embryos injected with either *Md-tra*-RNAi or GFP-RNAi were raised on porcine feces because the small number of injected larvae are less likely to develop into adult flies on standard rearing media (Schmidt *et al*. 1997). Under the injection scheme (Table 1), we could collect four types of flies: (A) genotypic females with the GFP-RNAi treatment (phenotypic females), (B) genotypic females with the *Md-tra*-RNAi treatment (sex-reversed males), (C) genotypic males with GFP-RNAi treatment (III^M^ males #1), and (D) genotypic males with the *Md-tra*-RNAi treatment (III^M^ males #2). Both types of genotypic males (III^M^ males #1 and #2) are also phenotypic males, and the GFP-RNAi treated genotypic females are phenotypic females. Sex reversal to a phenotypic male occurs in genotypic females under the *Md-tra*-RNAi treatment (Hediger *et al*. 2010).

**Table 1.** Injection scheme for RNAi treatments in both sexes.

| Genotypic Sex | RNAi treatment |  |
| --- | --- | --- |
|  | GFP-RNAi | *Md-tra*-RNAi |
| Genotypic Female<br>(III/III) | (A) Phenotypic Female<br>(III/III)<br>“females” | (B) Phenotypic Male<br>(III/III)<br>“sex-reversed males” |
| Genotypic Male<br>(III<sup>M</sup>/III) | (C) Phenotypic Male<br>(III<sup>M</sup>/III)<br>“III<sup>M</sup> males #1” | (D) Phenotypic Male<br>(III<sup>M</sup>/III)<br>“III<sup>M</sup> males #2” |

Genotypic females with the *Md-tra*-RNAi treatment are sex-reversed to phenotypic males (**B**). The other genotypic females and males are not sex-reversed (**A**, **C**, **D**); their phenotypic sexes are congruent with their genotypic sexes (i.e., normal males or females).

After emergence from pupa, each injected single phenotypic male was kept in a small cage with three or four females from the *pw bwb* strain that did not have any injection treatments. Only phenotypic males that successfully sired offspring with those females were retained for the RNA-seq experiment. All three types of phenotypic males produced offspring, but the sex-reversed males sired only female offspring (because they do not carry *Mdmd*). To measure gene expression in females, we collected virgin GFP-RNAi treated genotypic females. Those females were collected within 8 hours of emergence and kept separate from males to ensure they were virgin. The females were aged for five days, and we selected three females to dissect for RNA-seq experiments. We measured gene expression in virgin females to exclude mating effects on female gene expression.

### RNA-seq experiments

We used RNA-seq to measure gene expression in heads and abdomens from individual males of the four strains carrying either the Y^M^ or III^M^ proto-Y chromosomes (Figure 1). The larvae were raised at 25°C on a standard diet of wheat bran, calf manna, yeast, reptile litter, and water, as described previously (Hamm *et al*. 2009; Meisel *et al*. 2015). Unmated adult males and females were sorted within 8 hours of emergence, kept separately at 22°C, and provided water, sugar, and powdered milk *ad libitum*. Heads and abdomens from adult flies at five days post emergence were dissected and frozen at −80°C. The heads and abdomens from individual males were homogenized in TRIzol Reagent, and then RNA was extracted using the Zymo Direct-zol kit following the manufacturer’s protocol including DNA digestion steps. Three biological replicates (i.e., three individual male heads and abdomens) were prepared from males of each of the four strains. Because females of three strains (CS, CSrab, and IsoCS) are isogenic, we sampled only one female from each of the strains. However, the RNA-seq library preparation for CS female abdomen failed, so that the female abdomen had only two biological replicates.

We also performed RNA-seq on heads and abdomens from the four types of flies injected with dsRNA (Table 1). Individual four to five-day old adult flies (described above) were frozen in liquid nitrogen, and RNA was extracted from the individual flies with the NucleoSpin RNAII kit (Macherey-Nagel, Düren, Germany) following the protocol of the manufacturer, which includes DNA digestion steps. Three biological replicates (i.e., three individual flies) from each of the four genotype-by-treatment combinations were collected.

RNA-seq libraries were prepared with the Illumina TruSeq Stranded mRNA Sample Preparation Kit following the protocols of the manufacturer. The libraries were run in six lanes for 75 cycles (i.e., 75 nucleotide reads) on an Illumina NextSeq500 machine at University of Houston Seq-N-Edit Core. For the strains with different naturally occurring proto-Y chromosomes (Figure 1), two of three lanes contained ten libraries comprised of one replicate from each strain, sex, and body part (four strains of males plus one of the strains of females by two tissues): CS male head and abdomen, CSrab male head and abdomen, IsoCS male head and abdomen, bwbCS male head and abdomen, and female head and abdomen. The third lane contained nine of the samples described above, but no CS female abdomen because that library preparation failed. For the RNAi experiment we ran three lanes, and each lane contained eight library samples, one replicate from each genotype-by-treatment combination and body part: *Md-tra*-RNAi genotypic female head and abdomen (sex-reversed male), *Md-tra*-RNAi genotypic male head and abdomen, GFP-RNAi genotypic female head and abdomen, and GFP-RNAi genotypic male head and abdomen.

### Data analysis

Illumina RNA-seq reads were aligned to house fly genome assembly v2.0.2 and annotation release 102 (Scott *et al*. 2014) using HISAT2 v2.0.1 (Kim *et al*. 2015). Read mapping information for the strains with different naturally occurring proto-Y chromosomes and flies from the RNAi experiment are shown in Supplementary Tables 1 and 2. Read coverage across the sex determining genes *Md-tra*, *doublesex* (*Md-dsx*), and *fruitless* (*Md-fru*) was determined with the ‘mpileup’ function in SAMtools (Li *et al*. 2009). Second, the aligned reads were assigned to all annotated genes with htseq-count in HTSeq v0.9.1 (Anders *et al*. 2015), with the --stranded=reverse option because we generated stranded RNA-seq libraries.

The HTSeq output was used as input into DESeq2 v1.16.1 to identify differentially expressed genes (Love *et al*. 2014). For the DESeq2 analysis of the four strains with different naturally occurring Y^M^ and III^M^ chromosomes, we performed pair-wise comparisons between males of each strain. We also performed pair-wise comparisons of males from each strain against females. For the RNAi experiment, we created a model in DESeq2 in which gene expression is predicted by genotypic sex, RNAi treatment (GFP-RNAi or *Md-tra*-RNAi), and the interaction between genotypic sex and RNAi treatment. The model allows for pair-wise comparisons between individuals with either the same genotypic sex or RNAi treatment. From the pair-wise comparisons, log_2_ fold-changes (log_2_FC) were extracted for each gene with false discovery rate corrected *P* values (Benjamini and Hochberg 1995). We also extracted log_2_FC for III^M^ males #2 over females using the equation: log_2_(III^M^ males #2 / females) = log_2_(III^M^ males #1 / females) + log_2_(III^M^ males #2 / III^M^ males #1). We cannot calculate a *P* value for a test of whether log_2_(III^M^ males #2 / females) is different from zero because it is not a pair-wise comparison performed by the model we created in DESeq2. Only genes with adjusted *P* values reported by DESeq2 are presented and used for downstream analyses. In other words, we considered a gene to be expressed if there was enough data to compare gene expression levels, and we ignored genes where a statistical test was not performed because expression was too low.

We performed a principal component (PC) analysis, used a grade of membership model implemented in the R package ‘CountClust’ (Dey *et al*. 2017), and performed hierarchical clustering to analyze the normalized expression count data from DESeq2. For the PC analysis (PCA) and hierarchical clustering, we used a regularized log transformation of the normalized count data calculated using the ‘rlog’ function in DESeq2 (Love *et al*. 2014). Because genes with low counts show the highest relative differences among samples and create large variances, these low count genes dominate the results of the PCA. The regularized log transformation stabilizes the variance of the data, making it homoscedastic. The hierarchical clustering was performed using the pheatmap() function in R (Kolde 2019). Gene Ontology (GO) terms were analyzed with DAVID v.6.8 (Huang *et al*. 2009).

We assigned house fly genes to chromosomes using the conservation of Muller elements across flies (Foster *et al*. 1981; Weller and Foster 1993), as done previously (Meisel *et al*. 2015; Meisel and Scott 2018). Briefly, the house fly and *Drosophila* genomes are organized into six chromosome arms (Muller elements A-F). Elements A-E correspond to the house fly chromosomes that were historically considered the autosomes. Element F is the historical house fly X chromosome (Vicoso and Bachtrog 2013). One-to-one orthologs between house fly and *Drosophila melanogaster* genes were identified as part of the house fly genome annotation (Scott *et al*. 2014). We assigned house fly scaffolds to Muller elements using a “majority rules” approach—if the majority of genes on a scaffold were orthologous to *D. melanogaster* genes on a single Muller element, then the house fly scaffold was assigned to that Muller element. In turn, all genes on that scaffold are assigned to the same Muller element.

### Data Availability

All RNA-seq data are available from the NCBI Sequence Read Archive within BioProject accessions PRJNA522991 (GEO accession GSE126685) and PRJNA522995 (GEO accession GSE126689).

## Results

### Different proto-Y chromosomes have minor effects on gene expression

We previously observed that hundreds of genes are differentially expressed between males carrying III^M^ and males carrying Y^M^ (Meisel *et al*. 2015). Those experiments controlled for genetic background by introgressing a Y^M^ chromosome onto the same background as the III^M^ chromosome. However, the previous work did not compare the effects of the proto-Y chromosome with the effect of introducing a new autosome on the common genetic background. It is therefore not clear if the expression differences between Y^M^ and III^M^ males were specific to introducing a new proto-Y chromosome onto a genetic background, or if changing any single chromosome can induce similar expression effects. To address this question, we used RNA-seq to measure gene expression in males from four nearly isogenic strains carrying either a Y^M^ or III^M^ chromosome (Figure 1). Two strains with “III^M^ males” have different III^M^ chromosomes on a common genetic background. A third strain with “Y^M^ males” has a Y^M^ chromosome, instead of III^M^, on the same genetic background. The fourth strain carries a different Y^M^ chromosome and a single copy of a different standard third chromosome (without *Mdmd*) on the same genetic background as the other three strains. If the III^M^ chromosome has a disproportionate effect on gene expression, we expect to observe more genes differentially expressed between III^M^ and Y^M^ males than between Y^M^ males that differ from each other by a single standard third chromosome.

To compare gene expression profiles across the strains, we used both a PCA and a grade of membership model (Dey *et al*. 2017). We excluded one of the three batches of male samples (one replicate from each of the four strains) in both abdomen and head because males in that batch had outlier expression profiles. Specifically, the expression profiles of males from the outlier batch did not cluster with the replicates with the same genotype from the other two batches in our PCA (Supplementary Figure 3). Similarly, in a grade of membership model, males from the outlier batch had dramatically different group assignments than males with the same genotype from the other two batches (Supplementary Figure 3).

After excluding the outlier batch, we find that sex explains much of the variance in gene expression in both abdomen and head. In abdomen, the first PC (PC1) and second PC (PC2) explain 84% and 7% of variance in gene expression across samples, respectively (Figure 2A). In head, PC1 and PC2 explain 43% and 28% of variance, respectively (Figure 2B). In both abdomen and head, all males from the four strains are separated from females along PC1. Notably, the two Y^M^ strains (that differ from each other by a single copy of a standard third chromosome) have the greatest separation of any pair of male samples along PC2 in the abdomen data. We observed similar results with a grade of membership model: males from all four strains show different membership from females in abdomen and head, and males from the two different Y^M^ strains have the most different membership composition (Supplementary Figure 3C). In head, one of the III^M^ genotypes is separated from the other males along PC2 (Figure 2B). In neither abdomen nor head do Y^M^ and III^M^ males have the greatest separation, suggesting that the Y^M^ and III^M^ chromosomes affect gene expression to a similar extent as a non-*Mdmd*-bearing, standard third chromosome.

**Figure 2.**
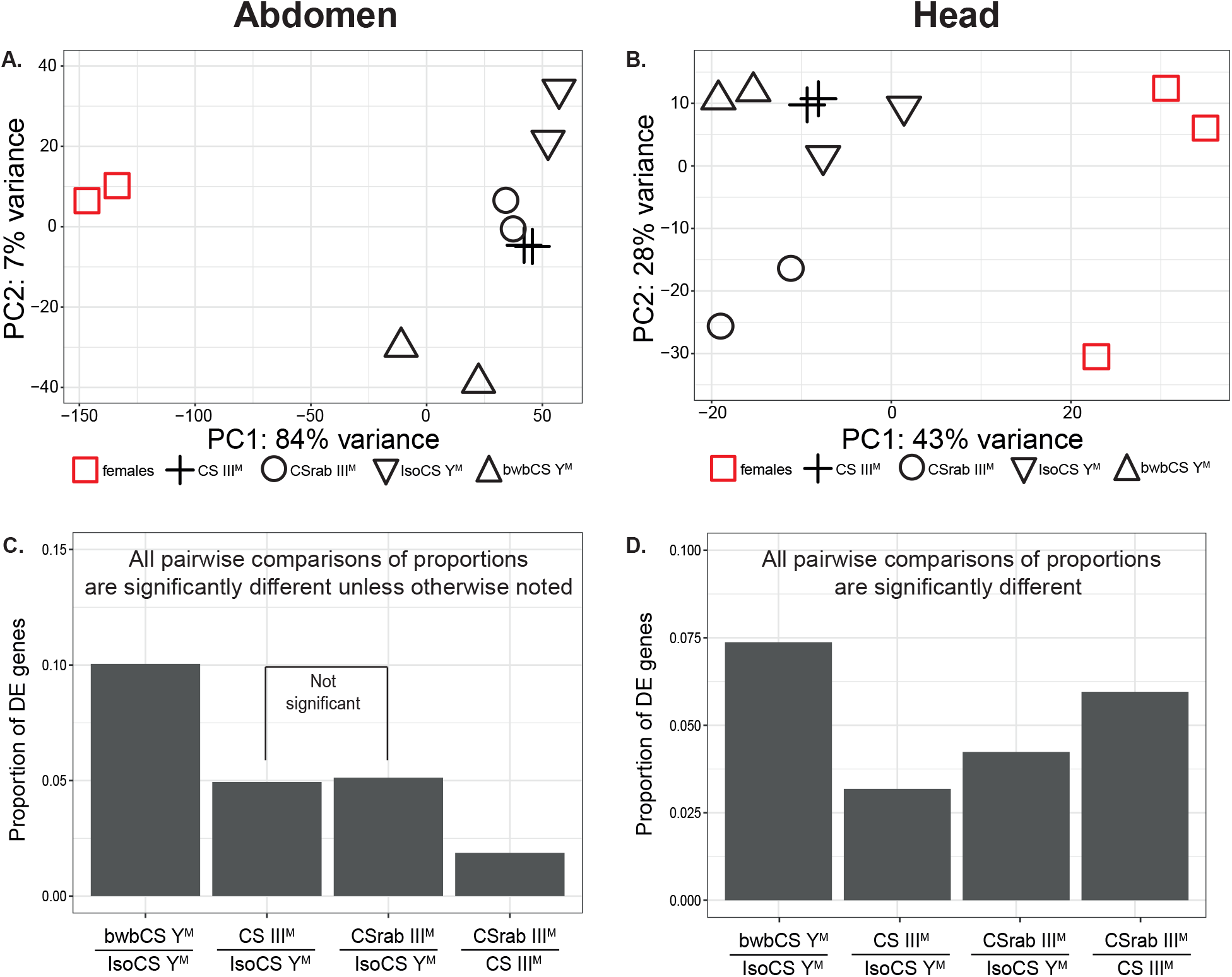
Plots show the results of a PCA of global expression in males with different proto-Y chromosomes in abdomen (**A**) and head (**B**). Bar graphs show the proportions of differentially expressed (DE) genes between males with different proto-Y chromosomes in abdomens (**C**) and heads (**D**). bwbCS Y^M^ stands for the strain bwbCS×CS.

We also identified individual genes with significant differential expression between strains using DESeq2 (Love *et al*. 2014). Previous work found that an excess of third chromosome genes is differentially expressed between Y^M^ and III^M^ males (Meisel *et al*. 2015), as expected based on the differences in their genotypes. We similarly find that excesses of genes on the third chromosome are differentially expressed in 5/8 comparisons between males with different III^M^ chromosomes, between Y^M^ males that differ by a standard third chromosome, and between Y^M^ and III^M^ males (Supplementary Figure 4). Only one other chromosome has a significant excess of differentially expressed genes in a single comparison (chromosome II in the bwbCS-IsoCS comparison). This amounts to one significant excess out of 32 total tests involving the other major chromosomes (chromosomes I, II, IV, and V are involved in 4 pair-wise comparisons from two tissue samples), which is less than the expected false-positive rate with a p-value cutoff of 0.05. Therefore, there is an excess of third chromosome genes with differential expression between males that differ in their third chromosome genotype, regardless of whether the third chromosome is a proto-Y (i.e., III^M^) or autosomal.

There are more differentially expressed genes across the entire genome in the pair-wise comparison between Y^M^ males with different standard third chromosomes than in any other pair-wise comparison between males, including between Y^M^ and III^M^ males (Figure 2C-D, Supplementary Figure 5, Supplementary Table 3). There are also similar fractions of differentially expressed (sex-biased) genes when comparing each of the male genotypes with females (Supplementary Figure 6). This provides further evidence that the III^M^ chromosome has a minor effect on male gene expression, less than or equal to an autosomal third chromosome. In summary, the PCA, grade of membership, and differential expression analyses all provide consistent evidence that the III^M^ chromosome has a minor effect on global gene expression.

### Expression of genes in the house fly sex determination pathway following *Md-tra* knock down

To further examine the effect of the III^M^ chromosome on gene expression, we used RNAi targeting *Md-tra* to create sex-reversed males that have a male phenotype and female genotype without any male-determining proto-Y chromosome. The *Md-tra*-RNAi treatment causes sex reversal by mimicking the effect of the male-determining *Mdmd* gene, which disrupts the splicing of *Md-tra* and the positive autoregulatory function of *Md-tra* in the early embryo (Hediger *et al*. 2010). We compared gene expression in the sex-reversed males with genotypic males carrying a III^M^ chromosome. Our 2×2 experimental design consisted of injecting dsRNA targeting either *Md-tra* (to sex-reverse genotypic females) or GFP (sham treatment) into genotypic males and females (Table 1). We studied III^M^ males in this experiment because we can use a morphological marker on the third chromosome to identify sex-reversed males (Hediger *et al*. 2010).

We confirmed that *Md-tra*-RNAi knockdown does indeed result in reduced expression of *Md-tra* in the sex-reversed male abdomens (Supplementary Figure 7A). In contrast, there is not a significant difference in *Md-tra* expression between sex-reversed male heads and either normal male or normal female heads (Supplementary Figure 7B). We discuss this observation further in Supplementary Figure 7. We observe sex-reversal in the expression of *Md-dsx* and *Md-fru*, the two immediate downstream targets of *Md-tra* (Hediger *et al*. 2004; Hediger *et al*. 2010; Meier *et al*. 2013), in *Md-tra*-RNAi treated genotypic females (Supplementary Figure 7C-F).

### Expression profiles of sex-reversed males are similar to genotypic males, not phenotypic females

We next examined how the III^M^ chromosome affects gene expression in males using a PCA analysis, grade of membership model, and hierarchical clustering to analyze RNA-seq data from the four genotype-by-RNAi-treatment combinations (Table 1). In abdomen, PC1 explains 85% of the variance in gene expression levels across samples. PC1 clusters all types of phenotypic males together, including the sex-reversed males, separately from normal females (Figure 3A). We observe a similar result in a grade of membership model and hierarchical clustering—the sex-reversed males have similar abdominal expression profiles as genotypic males and different from normal females (Supplementary Figures 8A and 9A). Therefore, the global gene expression profile in abdomen is predicted by phenotypic sex (regulated by the sex determination pathway), and not by sex chromosome genotype. This is consistent with the III^M^ chromosome having a minor effect on male gene expression.

**Figure 3.**
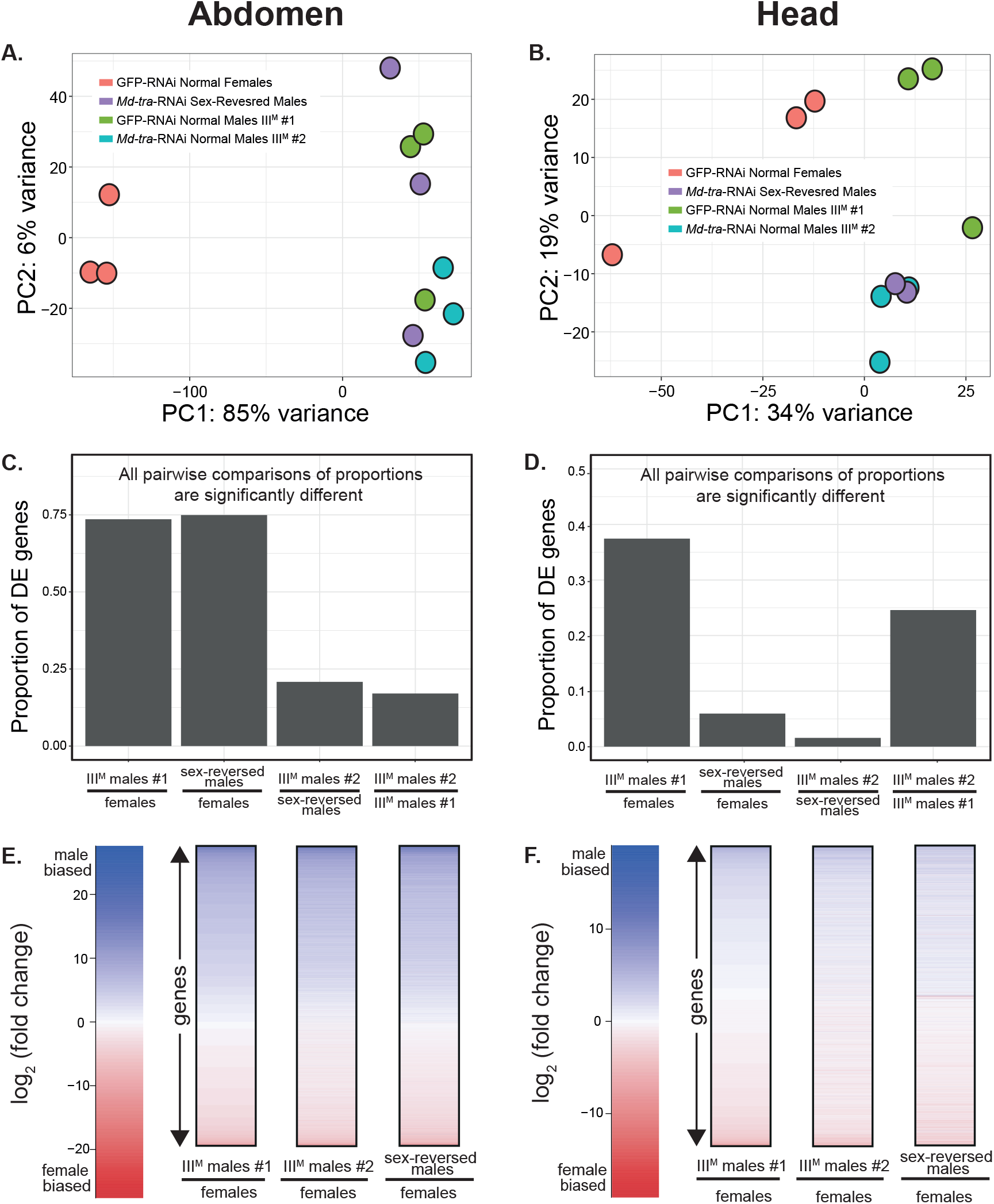
Plots show the results of a PCA of global expression of GFP-RNAi and *Md-tra*-RNAi treated genotypic females and males in abdomen (**A**) and head (**B**). Bar graphs show the proportions of differentially expressed (DE) genes between different types of individuals in abdomen (**C**) and head (**D**). “Females” refers to GFP-RNAi treated normal females. Heat maps show expression differences between each type of male and phenotypic females in abdomen (**E**) and head (**F**). Genes are ordered the same in all three comparisons within each tissue, such that a row in comparison contains the same gene as the corresponding row in the other two comparisons.

In head, we found that one of the sex-reversed males had elevated *Md-tra* expression (Supplementary Figure 10A), suggesting incomplete knock down of *Md-tra*. In addition, the RNA-seq profile of that sex-reversed male revealed it to be an outlier that did not cluster with either normal females or genotypic males in a PCA (Supplementary Figure 10B). The PCA showed that more variance in head gene expression can be explained by differences between that outlier sex-reversed male and all other phenotypic males (including the other two sex-reversed replicates) than between phenotypic males and females (Supplementary Figure 10B). We therefore excluded that outlier sample because it is not consistent with sex-reversal.

After excluding the one sample, PC1 and PC2 explain 34% and 19% of the variance in gene expression in head, respectively. PC1 for the head expression data separates normal females and GFP-RNAi treated genotypic males (Figure 3B), providing more evidence that phenotypic sex is a better predictor of overall gene expression profile than sex chromosome genotype. Curiously, *Md-tra*-RNAi treated phenotypic males (which includes both sex-reversed males and III^M^ males #2) were intermediate between GFP-RNAi treated normal females and males along head PC1, and the *Md-tra*-RNAi flies were separated from GFP-RNAi treated flies along head PC2 (Figure 3B). The *Md-tra*-RNAi treated phenotypic male heads (sex-reversed males and III^M^ males #2) also had reduced expression of *Md-fru* (Supplementary Figure 7F). We confirmed the similarity between *Md-tra*-RNAi treated flies, regardless of genotypic sex, in both a grade of membership model and via hierarchical clustering (Supplementary Figures 8B and 9B). Therefore, *Md-tra* knock down appears to influence overall gene expression—as well as *Md-fru* expression or splicing—in heads more than the effect of sex chromosome genotype.

### Sex-reversed and genotypic males have similar sex-biased gene expression

Sexual dimorphism is achieved through differential (sex-biased) gene expression between males and females (Ellegren and Parsch 2007). We compared sex-biased expression in sex-reversed and genotypic males, excluding the one outlier sex-reversed male head. We used genotypic males treated with GFP-RNAi (III^M^ males #1) as our normal male reference because the model in DESeq2 we used for RNA-seq analysis allows for pair-wise comparisons between individuals with either the same genotypic sex or treatment. Normal females and III^M^ males #1 both were exposed to the GFP-RNAi treatment, which allows us to make the pairwise comparison (Table 1).

We first quantified the degree of sex-biased expression by the distribution of the log_2_ fold-change between male and female expression levels (log_2_M/F). In the abdominal samples, the distributions of log_2_M/F for sex-reversed and genotypic males, when compared to normal females, are quite similar (Supplementary Figure 11A). We defined genes with sex-biased expression as those with a log_2_M/F significantly different from 0 using DESeq2 (Love *et al*. 2014). Similar fractions of genes have sex-biased expression in abdomen for sex-reversed and genotypic males: 11,005/14,686 (74.9%) of genes are significantly sex-biased in the comparison between sex-reversed males and females, and 11,030/14,993 (73.6%) of genes have sex-biased expression when comparing genotypic males and females (Figure 3C; Supplementary Table 4). The distributions of log_2_M/F are symmetrical, with similar fractions of genes with male- and female-biased expression for both sex-reversed and genotypic males (Supplementary Figure 11A). In contrast, the magnitude of differential gene expression is much smaller in comparisons between genotypic males than male-female comparisons (Figure 3C; Supplementary Figure 11A). Notably, the proportion of differentially expressed genes is similar between sex-reversed and genotypic males as between the two types of genotypic males (*Md-tra*-RNAi and GFP-RNAi treated), providing additional evidence that sex-reversed males have similar gene expression profiles as normal (genotypic) males (Figure 3C).

Sex-biased expression in fly heads is reduced relative to whole fly or gonad tissue (Goldman and Arbeitman 2007; Lebo *et al*. 2009; Meisel *et al*. 2012, 2015). In house fly heads, we only detect 5,077 sex-biased genes between genotypic males and normal females out of 13,558 expressed genes (Figure 3D; Supplementary Figure 11B; Supplementary Table 4). Similarly, there are only 735 sex-biased genes between sex-reversed males and normal females out of 12,360 expressed genes (Figure 3D; Supplementary Figure 11B; Supplementary Table 4). The lower number of sex-biased genes between sex-reversed males and normal females could be a result of decreased power because of a smaller sample size; only two replicate sex-reversed male heads were included because the third replicate had an outlier expression profile (see above). The alternative—that sex-reversed male heads are less sexual dimorphic than genotypic male heads—is unlikely because sex-reversed male head gene expression profiles are most similar to genotypic males in both a PCA and (Figure 3B; Supplementary Figure 9B). In addition, there are fewer genes differentially expressed in head between sex-reversed males and *Md-tra*-RNAi treated genotypic males (III^M^ males #2) than between the two types of genotypic males (Figure 3D). This result is consistent with the grouping of sex-reversed male and *Md-tra*-RNAi treated genotypic male head gene expression profiles in both a PCA and hierarchical clustering (Figure 3B; Supplementary Figure 9B), suggesting that gene expression in head is more affected by *Md-tra*-RNAi than by the III^M^ chromosome.

We next tested if the same genes have sex-biased expression in sex-reversed males and genotypic males (III^M^ males #1). In both abdomen and head, the majority of male-biased genes in genotypic males are also male-biased in sex-reversed males (Figure 3E-F). The same is true for female-biased genes. We assessed if the sex-biased genes in common between sex-reversed and genotypic males is greater than expected by chance with a permutation test. We determined a null distribution assuming that sex-biased genes in the sex-reversed and genotypic males are independent of each other from 1,000 random permutations of our data. The actual number of male- and female-biased genes in common between sex-reversed males and genotypic males is much greater than all values in the null distribution (Supplementary Figure 12). This result implies that sexual dimorphism is achieved by similar means in both sex-reversed males and genotypic males: silencing of *Md-tra*, independent of alleles on the III^M^ chromosome.

### Disproportionate differential expression of third chromosome genes

Although all phenotypic males, regardless of genotypic sex, showed very similar gene expression profiles, we identified some genes that are differentially expressed between genotypic males and sex-reversed males (Figure 3). These differentially expressed genes could reveal important phenotypic effects of the III^M^ proto-Y chromosome, which may be targets for selection to maintain both Y^M^ and III^M^ across house fly populations. We therefore further examined differential expression between sex-reversed and genotypic males to determine the effect of the III^M^ chromosome on individual genes. As expected based on their genotypic differences, there are significant excesses of third chromosome genes differentially expressed between genotypic III^M^ males and sex-reversed males in abdomen and head (Figure 4). There is also a significant excess of third chromosome genes differentially expressed between genotypic males and normal females in head (Figure 4B). In contrast, there is not an excess of third chromosome genes differentially expressed between normal females and sex-reversed males (Figure 4), who share the same genotype. These patterns are consistent with our previous work (Meisel *et al*. 2015) and other results presented here (Supplementary Figure 4) in which the third chromosome has an excess of differentially expressed genes between flies that differ in their third chromosome genotype. However, we surprisingly find that there are excesses of differentially expressed genes on the third chromosome in comparisons between III^M^ males with the *Md-tra*-RNAi and GFP-RNAi treatments (Figure 4). Therefore, in addition to the expected genotypic effects, dsRNA targeting *Md-tra* and/or GFP disproportionately affects the expression of genes on the house fly third chromosome.

**Figure 4.**
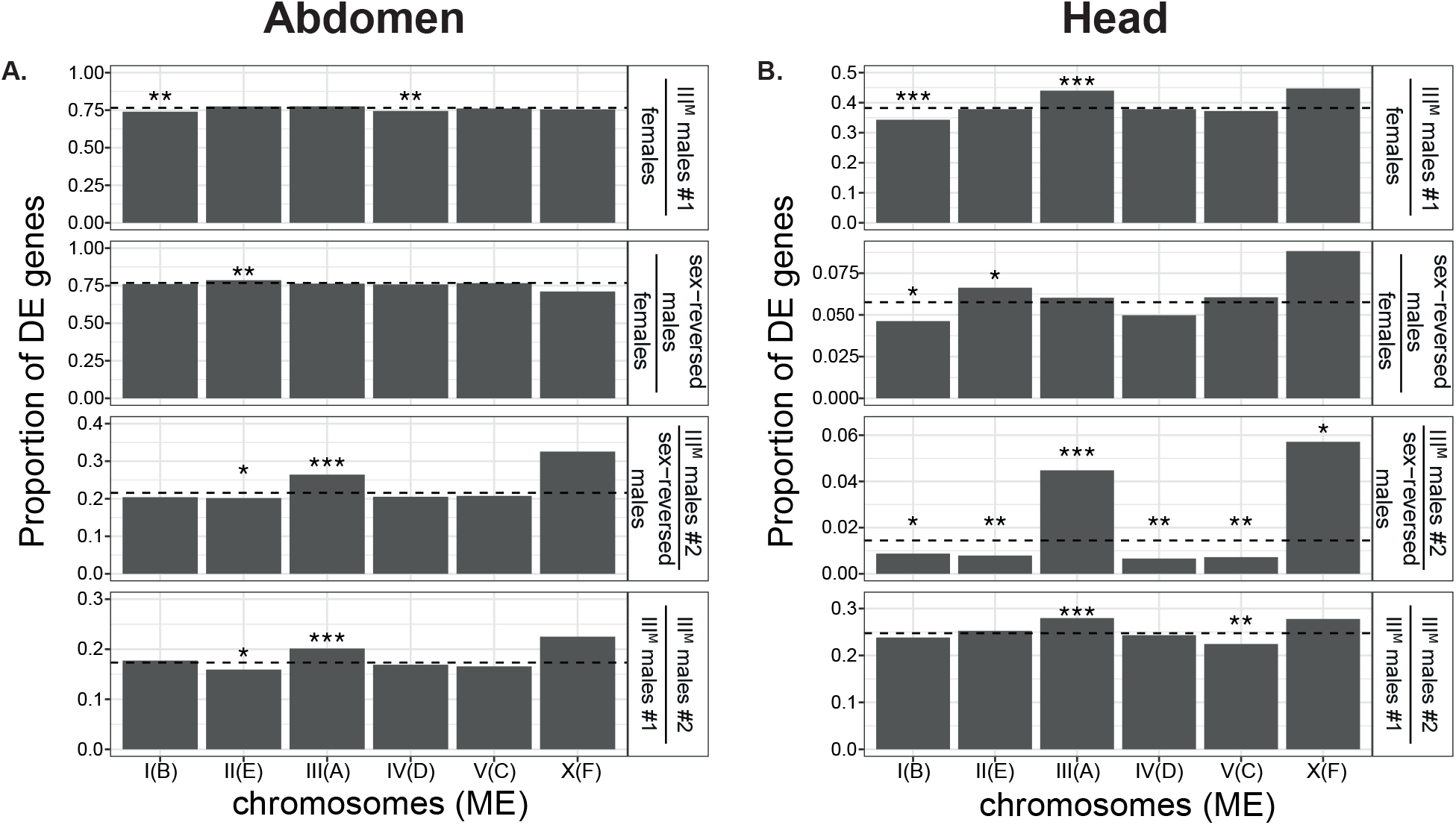
Bar graphs show the proportions of genes on each chromosome (*Drosophila* Muller element in parentheses) that are differentially expressed (DE) between genotype and treatment combinations in abdomen (**A**) and head (**B**). Asterisks indicate significant differences based on Fisher’s exact test (\**P*<0.05, \*\**P*<0.01, \*\*\**P* < 0.001).

Sex reversed males and genotypic males have very similar expression profiles (Figure 3). In spite of these similarities, we identified some “discordant sex-biased genes” that have sex-biased expression in either sex-reversed or genotypic males, but not both (Figure 3E-F). To further examine the effect of the III^M^ chromosome on gene expression, we divided the discordant sex-biased genes into two groups: “sex-reversed-up-discordant” and “normal-up-discordant”. We considered a gene to be sex-reversed-up-discordant if it belongs to one of two categories: 1) male-biased expression in sex-reversed males and not male-biased in genotypic males (log_2_M/F<0 but not necessarily significant), or 2) female-biased expression in genotypic males and not female-biased in sex-reversed males (log_2_M/F>0 but not necessarily significant). These genes are pooled into a single group because they have higher expression in sex-reversed males. We identified 49 sex-reversed-up-discordant genes in abdomens and 170 in heads (Figure 5A; Supplementary Table 5; Supplementary Data). Likewise, we classified genes as normal-up-discordant if they are in one of two categories: 1) male-biased expression in genotypic (normal) males and not male-biased in sex-reversed males (log_2_M/F<0 but not necessarily significant), or 2) female-biased expression in sex-reversed males and not female-biased expression in normal males (log_2_M/F>0 but not necessarily significant). These genes are pooled together because they have higher expression in normal males. We identified 25 normal-up-discordant genes in abdomens and 418 in heads (Figure 5B; Supplementary Table 5; Supplementary Data). There are no GO terms significantly enriched in either the sex-reversed-up-discordant or normal-up-discordant genes. However, both sex-reversed-up-discordant and normal-up-discordant genes in both abdomen and head are significantly enriched on the third chromosome (Figure 5; Supplementary Tables 6 and 7). Therefore, in comparisons between males with and without a III^M^ chromosome, the third chromosome is enriched for genes with discordant sex-biased expression.

**Figure 5.**
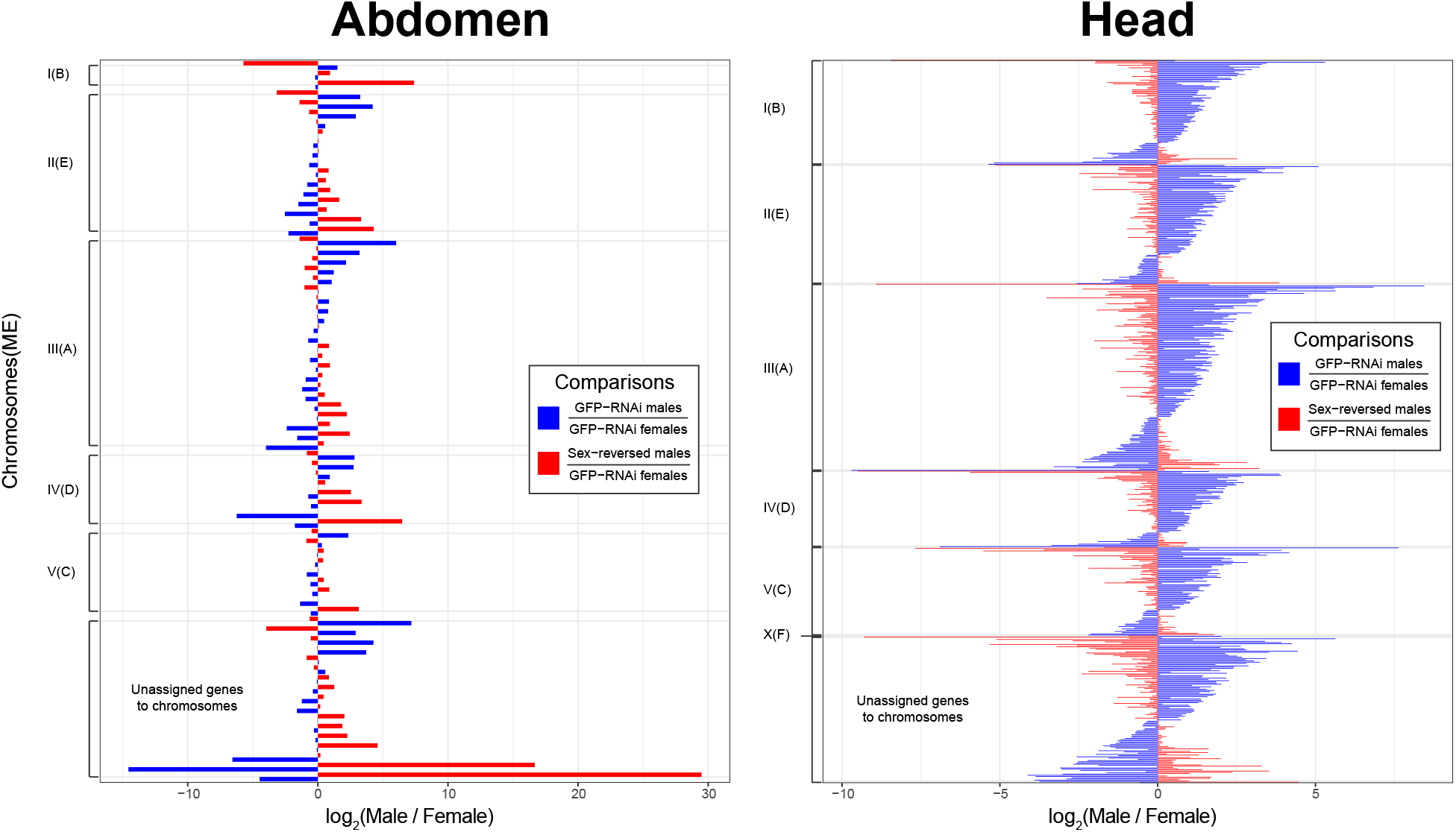
Chromosomal locations of discordant sex-biased genes in abdomen (**A**) and head (**B**) are shown. Blue bars indicate the degree of sex-biased expression (log_2_M/F) in genotypic males (III^M^ males #1) compared to females, and red bars show sex-biased expression in sex-reversed males compared to females. Each row represents a single gene with discordant sex-biased expression, with one blue bar and one red bar per gene.

## Discussion

The house fly Y^M^ and III^M^ proto-Y chromosomes are geographically distributed in a way that suggests ecological factors favor different proto-Y chromosomes across different habitats (Franco *et al*. 1982; Tomita and Wada 1989; Hamm *et al*. 2005; Feldmeyer *et al*. 2008; Kozielska *et al*. 2008). This predicts that there will be sequence differences between the proto-Y chromosomes and their homologous (proto-X) chromosomes that confer ecologically dependent phenotypic and fitness effects. These differences could be in transcribed sequences (e.g., protein coding genes) or in regulatory regions that control their expression. Paradoxically, however, both the Y^M^ and III^M^ chromosomes have minimal sequence differences relative to their homologous chromosomes (Meisel *et al*. 2017).

We tested if minimal sequence differences between the Y^M^ and III^M^ proto-Y chromosomes and their homologs could be responsible for phenotypic effects by investigating gene expression differences between males carrying different proto-Y chromosomes. We hypothesize that gene expression differences between males with different proto-Y chromosomes are caused by differences between proto-Y chromosomes and their homologs (e.g., III^M^ and the standard third chromosome) in their *cis*-regulatory sequences, the *trans*-regulatory factors they encode, and any downstream effects. The *cis*-regulatory differences should disproportionately affect expression of genes on the proto-Y chromosomes. Consistent with that prediction, there is a significant excess of genes that are differentially expressed on the third chromosome when comparing males with and without the III^M^ proto-Y (Figure 4; Supplementary Figure 4), as previously described (Meisel et al. 2015).

We compared gene expression in four house fly strains carrying either a Y^M^ or III^M^ chromosome on a common genetic background (Figure 1). The nearly isogenic strains were created by backcrossing the proto-Y chromosomes onto the common background (e.g., Supplementary Figure 1). Male recombination is absent in most flies (Gethman 1988), suggesting that our strains contain intact proto-Y chromosomes inherited from the progenitor strain. However, there is some evidence for male recombination in house flies (Feldmeyer et al. 2010). If recombination occurred between the proto-Y chromosomes and their homologs during backcrossing, then portions of the proto-sex chromosomes would also be isogenic across the strains we examined. Regardless of the extent of male recombination, expression differences between the strains should be (almost) entirely the result of genetic differences in proto-Y chromosome sequences (III^M^ or Y^M^) between strains, even if the entire proto-Y chromosome was not introgressed. Notably, the biggest differences in gene expression were observed between two Y^M^ strains carrying different standard third chromosomes (not carrying *Mdmd*), and not between III^M^ and Y^M^ males (Figure 2). Our results therefore suggest that the magnitude of gene expression differences between III^M^ and Y^M^ males can be explained by replacing a chromosome on a common genetic background, and they are not specific to the effect of the III^M^ or Y^M^ chromosomes.

We also examined the effects of the III^M^ chromosome on male gene expression using an RNAi experiment (Table 1). We chose to knock down *Md-tra* because it allows us to create sex-reversed fertile males that do not carry any proto-Y chromosomes, as opposed to knock down/out of *Mdmd*, which creates sex-reversed fertile females carrying a proto-Y (Sharma *et al*. 2017). We found that gene expression profiles of sex-reversed and normal (genotypic) males are very similar—phenotypic male expression profiles are distinct from phenotypic females, regardless of male genotype (Figures 3). We therefore conclude that the III^M^ chromosome has a minor effect on male gene expression in a constant environment as assayed in our experiments. It is worth noting that we only considered virgin females in our experiments, and mating can affect female gene expression in flies (e.g., McGraw et al. 2004; Dalton et al. 2010). However, it is unlikely that including mated females would have dramatically affected our central finding— mated females should have global gene expression profiles that are more similar to virgin females than phenotypic males.

Despite the minimal effect of the proto-Y chromosomes on global gene expression, we identified 74 genes in abdomen and 588 genes in head with sex-biased expression that is affected by the III^M^ chromosome. These genes with discordant sex-biased expression are enriched on the third chromosome (Figure 5). Discordant sex-biased genes could be targets of selection involved in the maintenance of polygenic sex determination. The small number of genes whose expression is affected by the proto-Y chromosomes suggests that the selection pressures responsible for maintaining the proto-Y chromosome polymorphism likely act on a limited number of genetic targets, not the entirety of the proto-Y chromosomes.

### Gene expression effects of the proto-Y chromosomes

A previous experiment identified many genes whose expression differs between Y^M^ and III^M^ males (Meisel *et al*. 2015). That experiment controlled for genetic background, but it did not compare the effect of the proto-Y chromosomes with the effects of equivalent autosomes. When we included the effect of introducing an autosome onto the same genetic background, we observe more expression differences between Y^M^ males that carry different copies of standard (non-*Mdmd*-bearing or autosomal) third chromosomes than between Y^M^ males and III^M^ males (Figure 2). This minimal effect of the III^M^ proto-Y chromosome on expression, relative to a standard third chromosome, suggests that III^M^ has very few differences from the standard third chromosome, and III^M^ is not a fully “masculinized” Y chromosome (Rice 1996). In addition, III^M^ likely is not differentiated enough from the standard third chromosome to require dosage compensation in heterogametic males. Alternatively, III^M^ males may have a dosage compensation mechanism (i.e., through preferred expression of genes on the standard third chromosomes), which could mask the effects of the III^M^ chromosome on gene expression. Additional work is necessary to compare these hypotheses.

It is curious that the Y^M^ males with different standard third chromosomes have more expression differences than between Y^M^ and III^M^ males (Figure 2). One explanation for the amount of expression differences between the Y^M^ males is that the standard third chromosome in our experiment has a greater effect on gene expression than the III^M^ chromosome. Alternatively, the different origins of the Y^M^ chromosomes in the two Y^M^ genotypes could have a large effect on gene expression. Unfortunately, our experimental design prevents us from differentiating between the effects of the Y^M^ chromosomes and standard third chromosome on the expression differences between these Y^M^ males. However, if differences between Y^M^ chromosomes were responsible for the elevated differential expression between the two Y^M^ male genotypes, this would suggest that variation amongst the effects of Y^M^ chromosomes in our experiment exceeds differences between Y^M^ and X chromosomes. Non-recombining Y chromosomes are expected to have low levels of polymorphism (Clark 1987, 1988). However, variation across *D. melanogaster* Y chromosomes has been shown to affect gene expression across the genome and may be involved in the resolution of sexual conflicts (Lemos *et al*. 2008, 2010). In addition, human Y chromosomes harbor high levels of copy number variation of ampliconic genes (Ye *et al*. 2018). Intriguingly, the house fly Y^M^ chromosome carries recently duplicated genes that differentiate it from the homologous X chromosome (Meisel *et al*. 2017). If these Y^M^ duplications vary in their copy number or if there are chromatin-level differences across Y^M^ chromosomes, this could explain a possible effect of the Y^M^ chromosome on global gene expression. Additional work is necessary to test these hypotheses.

### The III^M^ chromosome, *cis*-regulation, and sexual conflicts

Despite the minimal effects of the III^M^ chromosome on global gene expression profiles, we do identify two notable patterns across all types of males. First, higher proportions of genes on the third chromosome, relative to other chromosomes, are differentially expressed in many of our comparisons between males with different genotypes (Figure 4; Supplementary Figure 4). This might be indicative of divergence of *cis*-regulatory alleles between the III^M^ and standard third chromosomes. The differentially expressed third chromosome genes could have important phenotypic effects that could partially be responsible for fitness differences between males with and without the III^M^ chromosome. Those fitness differences could in turn explain the maintenance of both the Y^M^ and III^M^ proto-Y chromosomes in natural populations. Additional work is necessary to connect individual differentially expressed genes to fitness effects of the III^M^ chromosome.

Second, genes with discordant sex-biased expression between genotypic males and sex-reversed males are over-represented on the third chromosome (Figure 5). This is consistent with our previous results showing that the III^M^ chromosome disproportionately promotes male-biased expression (Meisel *et al*. 2015). These results are contingent on inference of the chromosomal assignment of house fly genes, which we have confirmed is accurate by comparing with an independent mapping approach (Meisel and Scott 2018).

Population genetics theory predicts that sexually antagonistic selection is a major driver of the evolution of sex determination and the maintenance of polygenic sex determination (Orzack *et al*. 1980; van Doorn and Kirkpatrick 2007, 2010; Meisel *et al*. 2016). For example, sexual conflicts could be resolved if sexually antagonistic alleles are inherited in a sex-limited manner through the origination of a tightly linked sex-determining factor (Lindholm and Breden 2002; Roberts *et al*. 2009; Ser *et al*. 2010; Parnell *et al*. 2013). In addition, male-beneficial alleles are expected to accumulate on proto-Y chromosomes once they have acquired male-limited inheritance (Rice 1992). The excess of discordant sex-biased genes on the third chromosome may be consistent with these theoretical predictions if the up- or down-regulation of these genes on the III^M^ chromosome is beneficial to males and deleterious to females. In this case, the male-beneficial (and female-detrimental) alleles would be *cis*-regulatory elements that affect the expression of the discordant sex-biased genes on the III^M^ chromosome (Figure 5). A similar phenomenon was observed in Lake Malawi cichlids, where an allele underlying a sexually antagonistic pigmentation phenotype is a *cis*-regulatory variant that up-regulates the expression of a gene linked to a new sex determiner (Roberts *et al*. 2009). Although the house fly male determiner (*Mdmd*) is molecularly characterized (Sharma *et al*. 2017), its location on the III^M^ chromosome is not known, which prevents us from testing if the discordant sex-biased genes are nearby and genetically linked to the male determiner.

There are two considerations, however, that may be important limitations of these interpretations. First, the fitness effects of the proto-Y chromosomes appear to be environmentally dependent. Y^M^ is most frequent at northern latitudes and III^M^ predominates in the south, suggesting that temperature-dependent fitness differences could be responsible for north-south clines (Franco *et al*. 1982; Tomita and Wada 1989; Hamm *et al*. 2005; Feldmeyer *et al*. 2008; Kozielska *et al*. 2008). We did not test for temperature-dependent effects of the proto-Y chromosomes in our experiment, which may have prevented us from identifying key fitness-related gene expression differences between Y^M^ and III^M^ males. These temperature-dependent effects could be the result of temperature-dependent expression of genes on Y^M^ and III^M^, differences in temperature-dependent activity of the copies of *Mdmd* across proto-Y chromosomes, or some other temperature-dependent genotype-by-environment interaction. Second, Y^M^ and III^M^ can be carried by females who also carry the epistatic *Md-tra^D^* allele (McDonald *et al*. 1978; Hediger *et al*. 2010). The fitness differences between Y^M^ and III^M^ could therefore be mediated through the effects of the proto-Y chromosomes on female phenotypes, which we did not assay in our experiments. Additional work is necessary to investigate how temperature and sex modulate the phenotypic effects of the proto-Y chromosomes.

### The effect of *Md-tra* on gene expression

Our results suggest that *Md-tra* has effects on gene expression beyond the direct regulation of *Md-dsx* and *Md-fru* splicing. Previous results, as well as our experiments here, demonstrate that knock down of *Md-tra* in blastoderm embryos causes complete sex-reversal of genotypic females into fertile phenotypic males (Hediger *et al*. 2010). Our results suggest that this sex-reversal does not affect all adult tissues equally—we observed one fertile sex-reversed male with higher *Md-tra* expression than normal females in head and a head gene expression profile that does not cluster with phenotypic females or males (Supplementary Figure 10). Curiously, the outlier sex-reversed male in our experiment does not have a gene expression profile intermediate between genotypic males and females (Supplementary Figure 10B), as we would expect from partial masculinization. This suggests *Md-tra*-RNAi treatment in blastoderm embryo can have effects on adult somatic gene expression that does not act in the expected direction of sex-reversal.

We find additional evidence that *Md-tra* knockdown can affect adult gene expression independently of genotype. For example, the two genotypic males in our RNAi knockdown experiment have the same genotypic and phenotypic sex, yet their head gene expression profiles are not the most similar of all genotype-by-treatment combinations; instead, genotypic males with *Md-tra*-RNAi treatment cluster with sex-reversed males (Figure 3B; Supplementary Figure 9B). There are also more genes differentially expressed in head between III^M^ males with and without *Md-tra*-RNAi treatment than between genotypic males and sex-reversed males (Figure 3D). These results suggest that *Md-tra* affects head gene expression independent of genotypic sex. The effects of *Md-tra*-RNAi on head expression are likely mediated through either direct effects of *Md-tra* on the splicing of transcripts other than *Md-dsx* and *Md-fru*, downstream effects of *Md-dsx* and *Md-fru* alternative splicing, or off-target effects of dsRNA targeting *Md-tra*. In contrast, we do not observe a disproportionate effect of *Md-tra*-RNAi on abdominal gene expression—knocking down *Md-tra* converts genotypic females into phenotypic (sex-reversed) males with expression profiles that nearly perfectly mimic genotypic (normal) males (Figure 3A,C,E; Supplementary Figures 8A and 9A).

Notably, the expression of *Md-tra* does not differ across the heads of genotypic males or females with either RNAi treatment (Supplementary Figure 7B). This suggests that the expression effects of knocking down *Md-tra* in adult heads is not through direct effects on *Md-tra*, but instead is caused by off-target effects or downstream effects of the direct targets of *Md-tra*. It is therefore possible that silencing *Md-tra* in early blastoderm embryos affects regulatory pathways that modulate head gene expression independently of the activity *Md-tra* in adult heads. Sex determination in flies is cell autonomous, and many cells in *Drosophila* somatic tissues do not express sex-determining genes downstream of *tra* (Robinett *et al*. 2010). Our results suggest that even if somatic tissues do not differentially express sex-determining genes, they can carry the memory of regulation of the sex determination pathway from their progenitor cells.

Curiously, there is an excess of third chromosome genes differentially expressed between III^M^ males with *Md-tra*-RNAi treatment and III^M^ males with *GFP*-RNAi treatment (Figure 4). The III^M^ chromosome is a proto-Y, and the standard third chromosome is a proto-X. Therefore, knockdown of *Md-tra* could be disproportionately affecting proto-Y genes or proto-X genes. Unfortunately, our data lack the resolution to determine if the expression changes between III^M^ males with different RNAi treatments is the result of changes in expression of genes on the III^M^ chromosome, standard third chromosome, or both. Regardless of which homolog is changing in expression, one explanation for the disproportionate effect of *Md-tra* knockdown on third chromosome genes is that there is an excess of third chromosome targets regulated by *Md-tra* or the sex determination pathway. For example, the house fly sex determination pathway could regulate gene expression specifically on the proto-X chromosome, analogous to how *Drosophila* X chromosome dosage compensation is controlled in a sex-specific manner by a gene in the sex determination pathway (Salz and Erickson 2010). Intriguingly, knockdown of *transformer* in female red flour beetles, *Tribolium castaneum*, causes them to produce nearly all male progeny, possibly as a result of misregulation of the diploid X chromosome in the female progeny (Shukla and Palli 2012). *Md-tra* in house fly may have a similar role regulating X chromosome expression. Additional work is necessary to evaluate why *Md-tra* knockdown disproportionately affects third chromosome expression.

## Conclusions

We have performed multiple RNA-seq experiments to resolve the paradox of ecologically relevant fitness effects of the house fly Y^M^ and III^M^ proto-Y chromosomes despite minimal sequence divergence between proto-Y and proto-X chromosomes. We identified some effects of the Y^M^ and III^M^ chromosomes on gene expression, but the number of differentially expressed genes and their effect sizes are small relative to the effect of a standard third chromosome or knockdown of the key sex determining gene *Md-tra*. Therefore, gene expression in house flies depends more on phenotypic sex (mediated by the sex determination pathway) than sex chromosome genotype. Despite the minimal effect of house fly proto-Y chromosomes on global gene expression, there are individual genes on the III^M^ chromosome that are differentially expressed relative to the standard third chromosome. Those differentially expressed genes may represent targets of selection that is responsible for maintaining polygenic sex determination.

Our results are consistent with a recent study in *Rana temporaria* frogs that have polygenic sex determination, which found that sex-biased gene expression depends more on phenotypic sex than genotypic sex (Ma *et al*. 2018). Combining the findings from flies and frogs suggests that the earliest stage of sex chromosome evolution is dominated by changes in the expression of individual genes, rather than transcriptome-wide effects of the young sex chromosomes. In the case of house fly, it is possible that the individual gene expression differences could act in an ecologically-dependent manner. For example, the geographic distribution of the Y^M^ and III^M^ chromosomes could arise from selection on environmentally sensitive phenotypes that we did not assay in our experiments. Because seasonality of temperature is predictive of the frequencies of Y^M^ and III^M^ in natural populations (Feldmeyer *et al*. 2008), a fitness or phenotypic assay across temperatures may be needed to identify ecologically relevant differences between Y^M^ and III^M^ males.

## Acknowledgements

This work was supported by a grant from the National Science Foundation (OISE 1444220) to RPM. We thank Christopher Gonzales for assistance with preparation of the RNA-seq libraries, and Claudia Brunner for technical assistance with the RNAi knockdown experiment. All RNA-seq data were generated at the University of Houston Seq-N-Edit core and analyzed on the Maxwell Cluster at the University of Houston Core facility for Advanced Computing and Data Science. We also thank Ernst Wimmer, Louis van de Zande, Elzemiek Geuverink, Martijn Schenkel, and Xuan Li for fruitful discussions on housefly sex determination evolution. Three anonymous reviewers provided valuable feedback that improved this manuscript.

## Supplementary Material

**Supplementary Figure 1.**
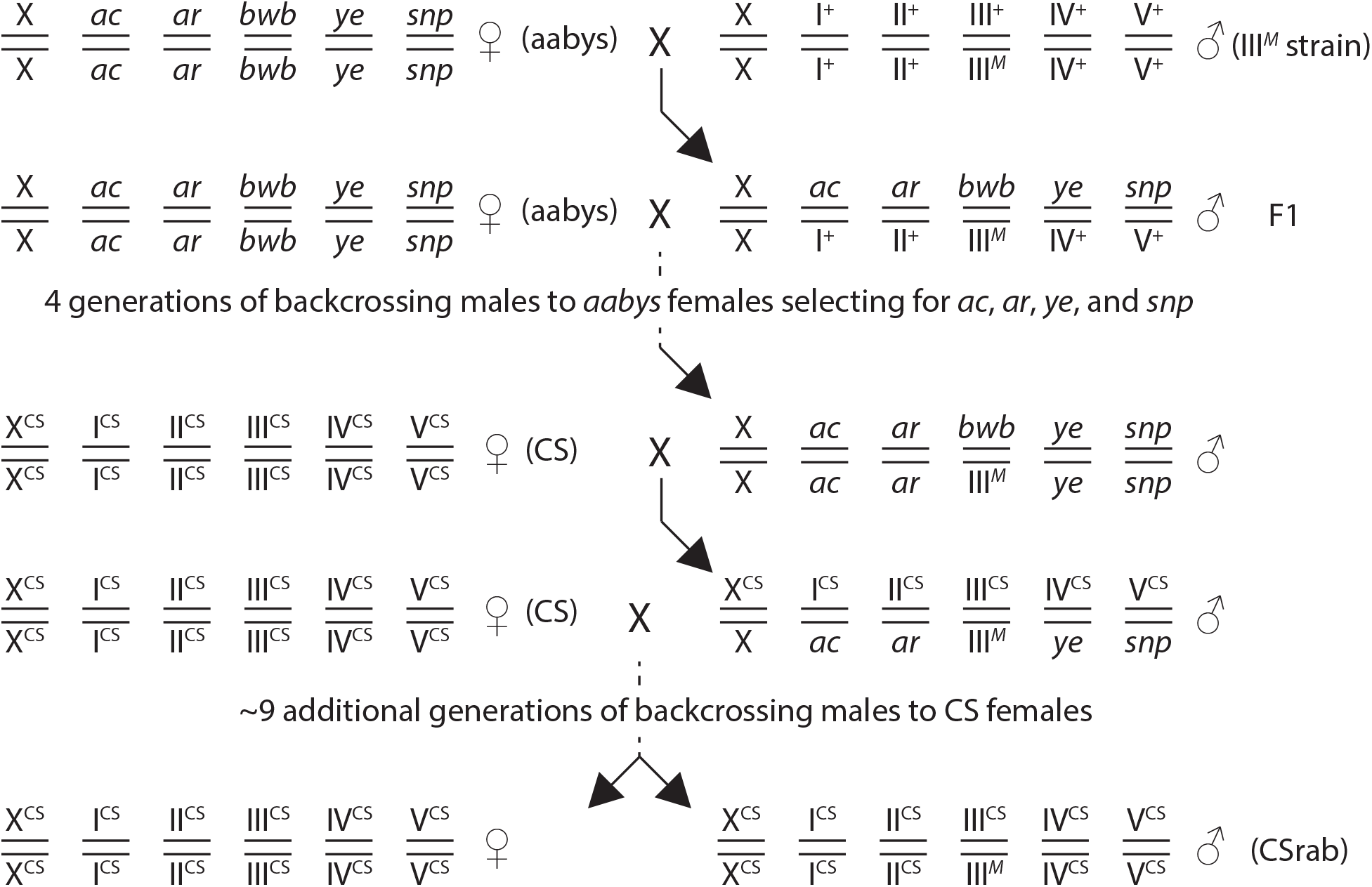
Crossing scheme used to create the CSrab strain. Each pair of parallel horizonal bars represents homologous chromosomes; there are six pairs of chromosomes (X/Y, I, II, III, IV, V). The aabys strain has a recessive phenotypic marker on each autosome.

**Supplementary Figure 2.**
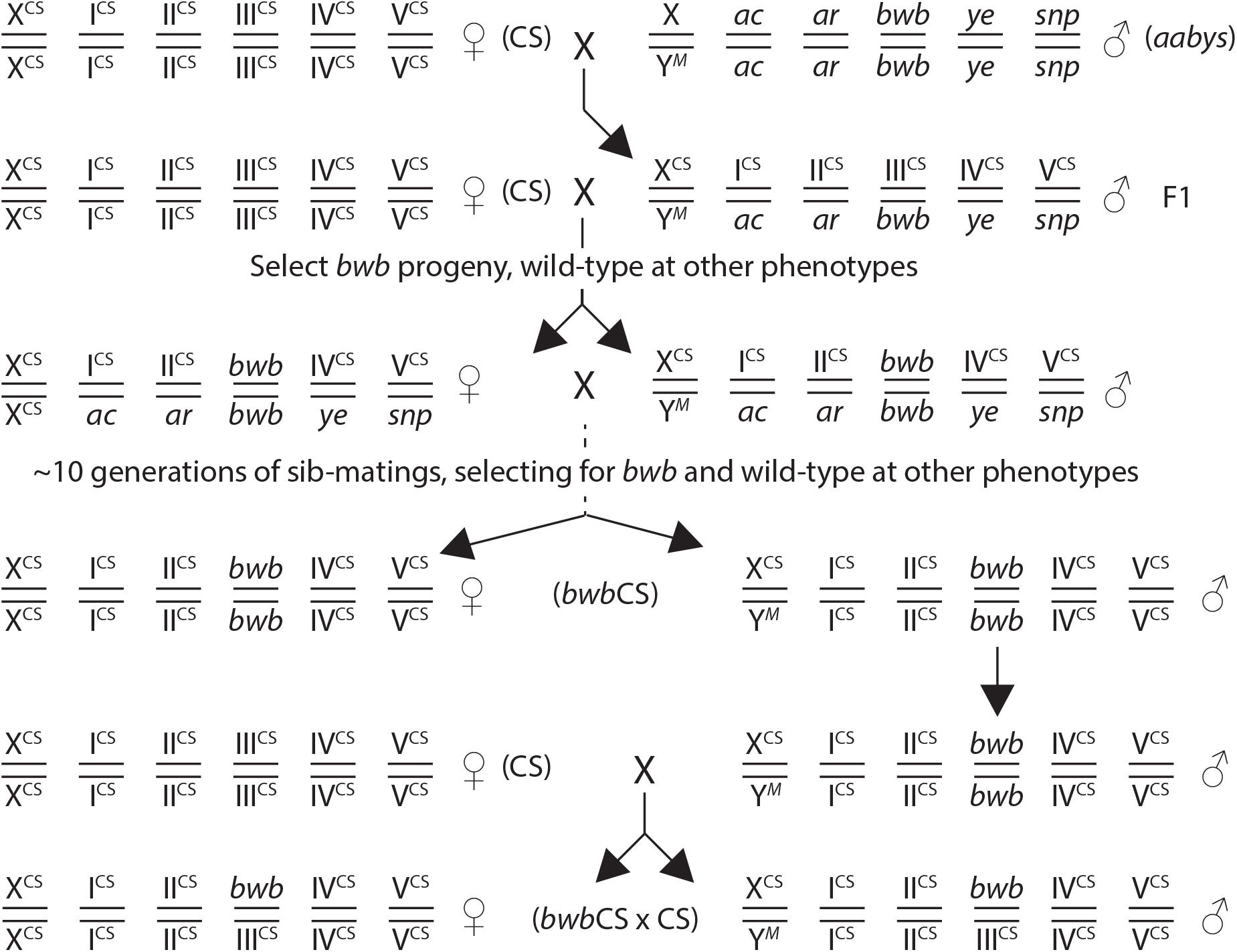
Crossing scheme to create the *bwb*CS strain and *bwb*CS × CS line used in this experiment. Each pair of parallel horizonal bars represents homologous chromosomes; there are six pairs of chromosomes (X/Y, I, II, III, IV, V). The aabys strain has a recessive phenotypic marker on each autosome.

**Supplementary Figure 3.**
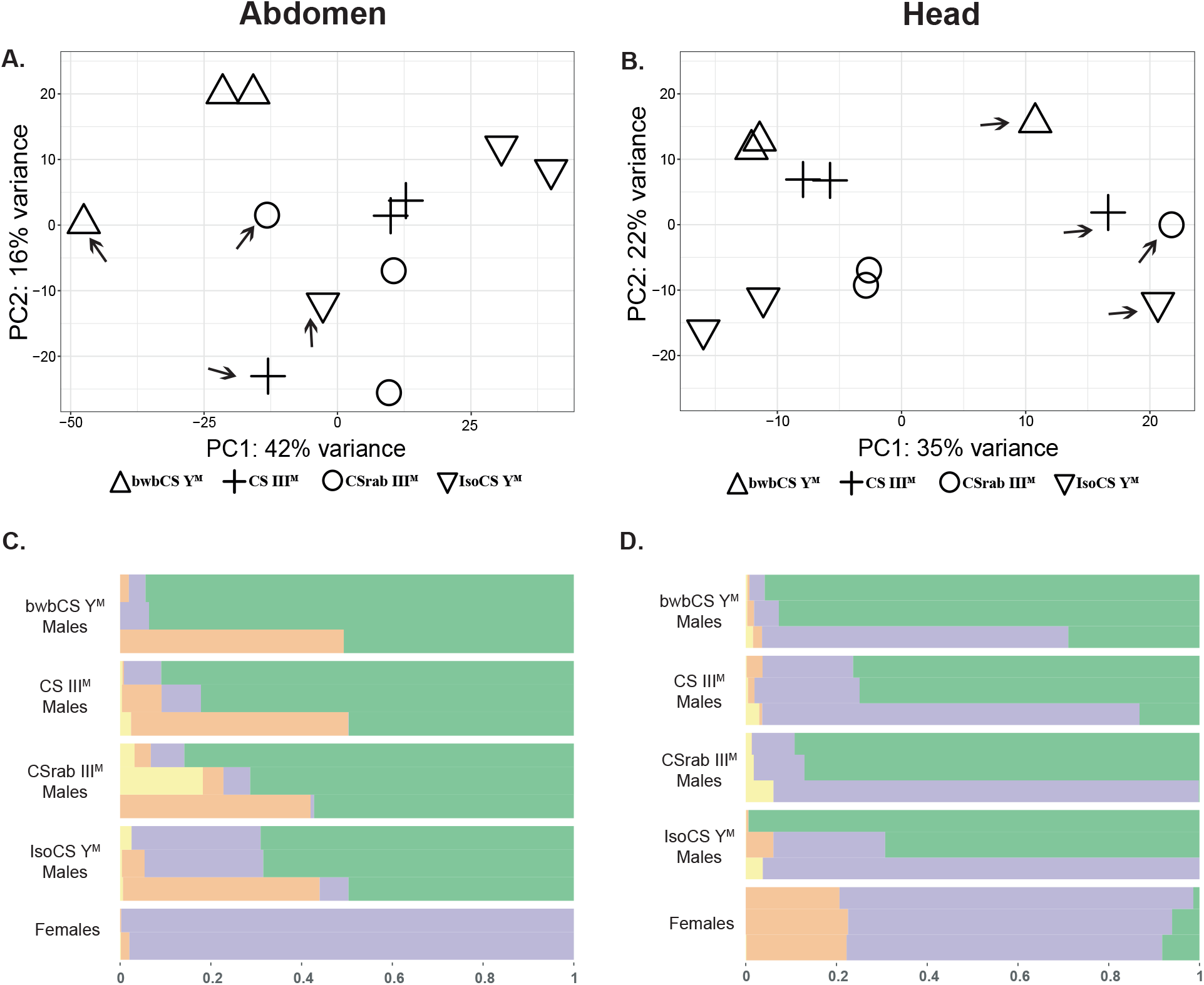
(**A, B**) Principal component (PC) analysis of four strains that have different naturally occurring proto-Y chromosomes on a common genetic background in abdomens (**A**) and heads (**B**). Arrows point to outlier samples, one for each of the four strains. Female abdomens are excluded from the PCA (**A**) to show the outliers. (**C, D**) Grade of membership model (*K* = 4) for gene expression patterns of four strains that have different naturally occurring proto-Y chromosomes on a common genetic background in abdomens (**C**) and heads (**D**). Each row represents one replicate of a genotype, with the outliers excluded. Each color represents the proportion of each replicate assigned to each of the three clusters.

**Supplementary Figure 4.**
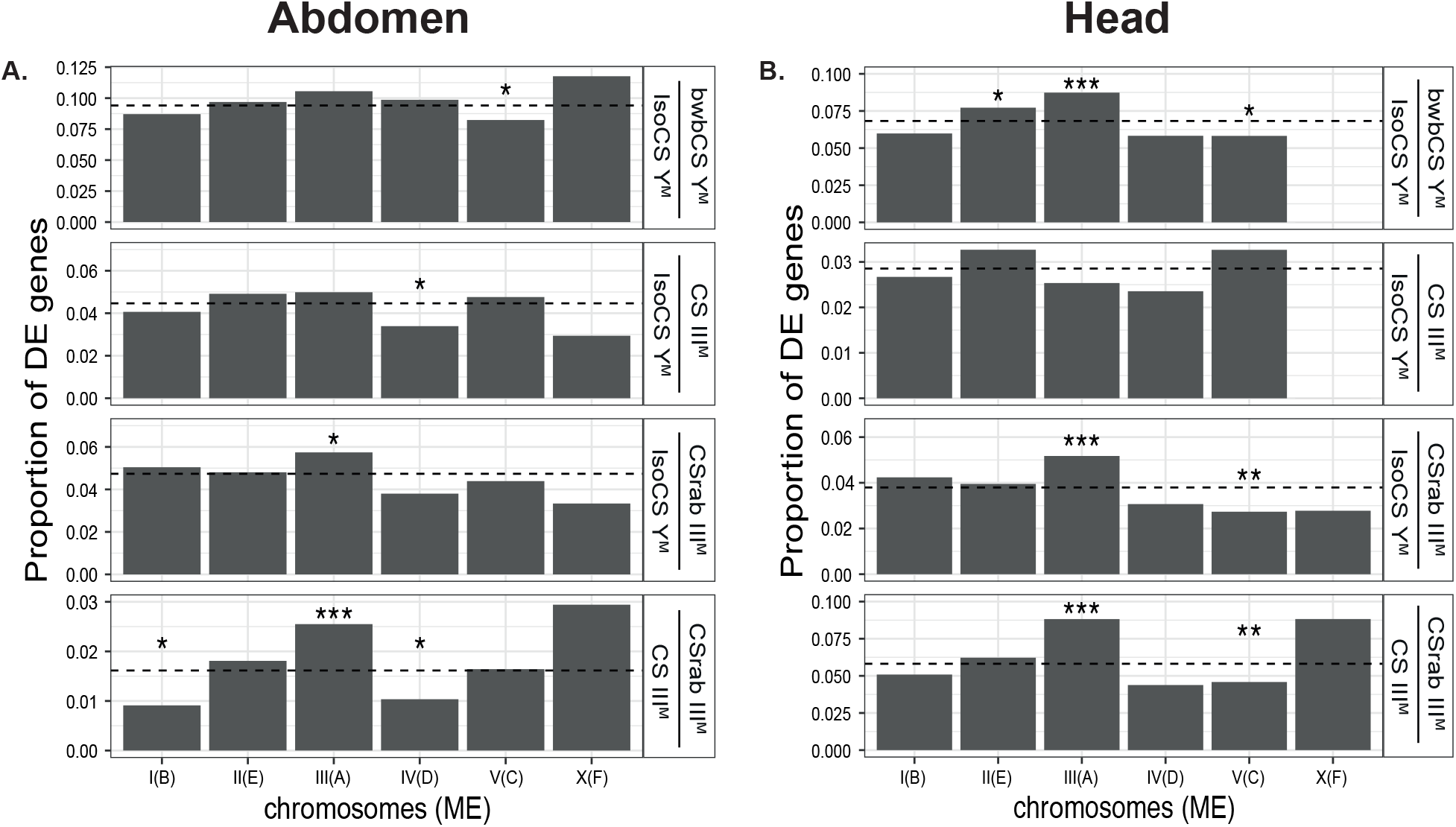
Bar graphs indicate the proportions of genes on each chromosome (*Drosophila* Muller element in parentheses) that are differentially expressed (DE) between different male genotypes in abdomen (**A**) and head (**B**). Asterisks indicate significant differences based on Fisher’s exact test comparing the number of DE genes on a chromosome against the number of DE genes in the rest of the genome (\**P*<0.05, \*\**P*<0.01, \*\*\**P* < 0.001).

**Supplementary Figure 5.**
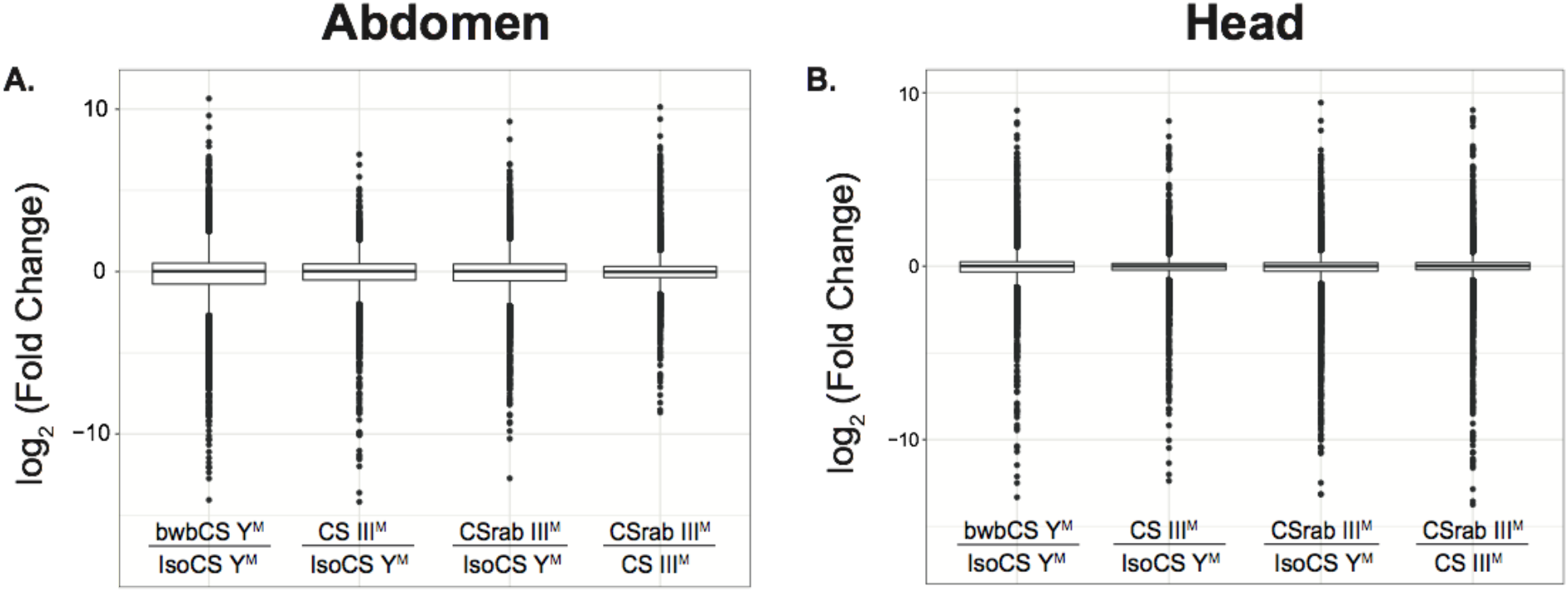
Boxplots show fold changes of gene expression between males with different *Mdmd*-bearing chromosomes in abdomens **(A)** and heads **(B)**. Outliers are included as points.

**Supplementary Figure 6.**
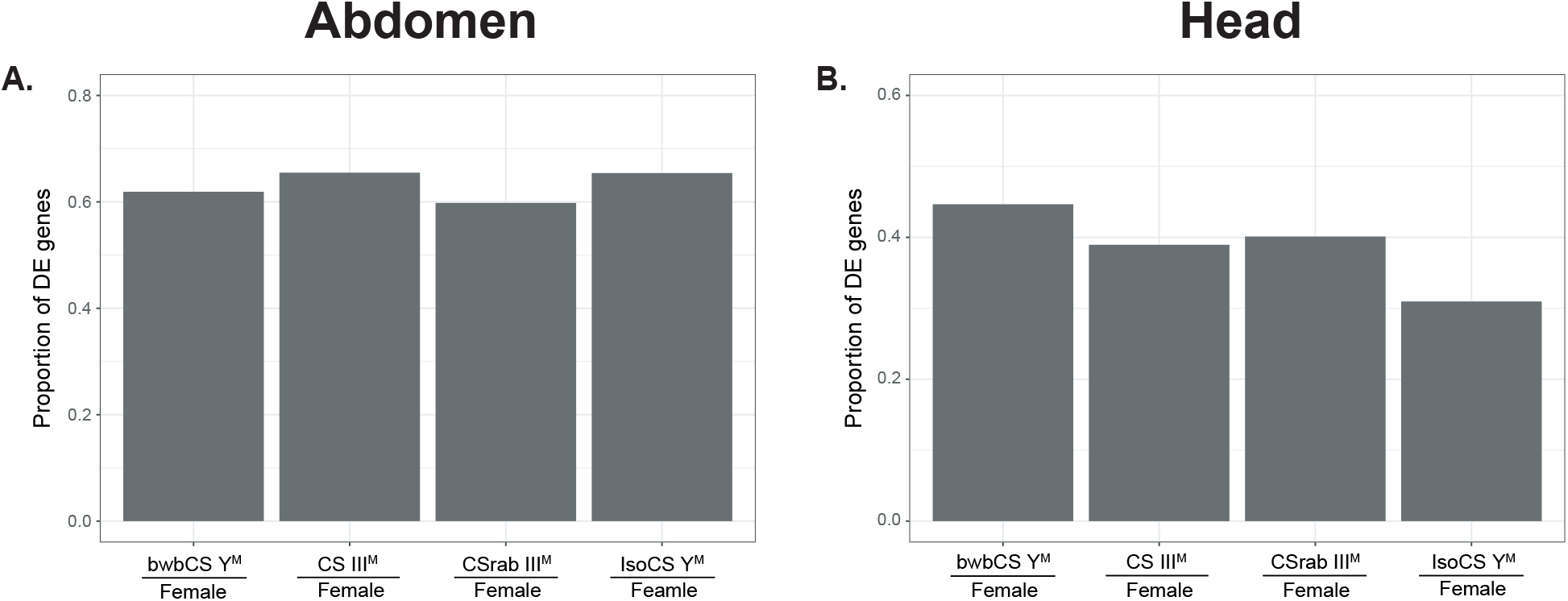
Bar graphs show the proportions of differentially expressed (DE) genes between females and males with different Y^M^ or III^M^ proto-Y chromosomes in abdomens (**A**) and heads (**B**). bwbCS Y^M^ stands for the strain bwbCS×CS.

**Supplementary Figure 7.**
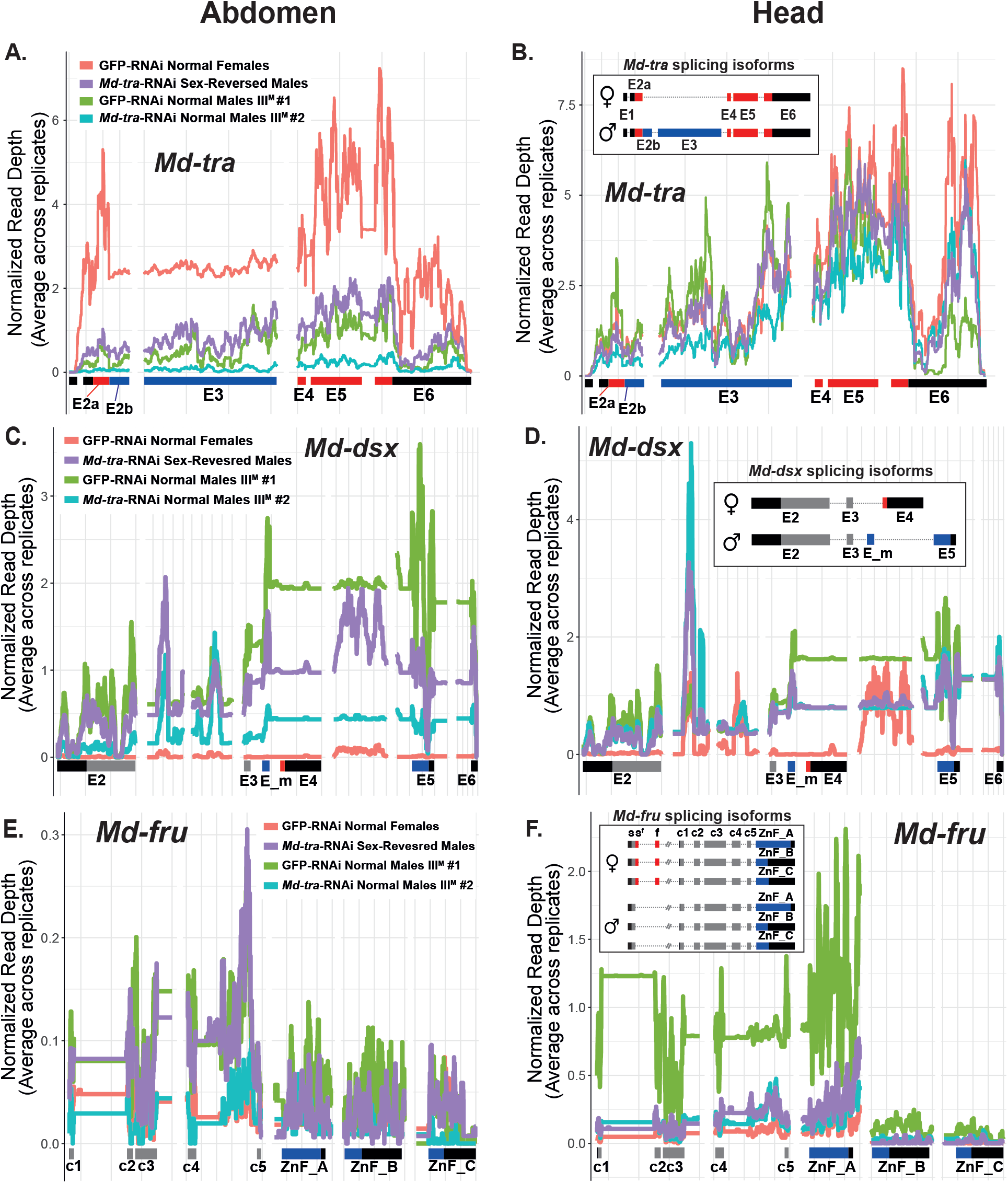
The graphs show read depth coverage of *Md-tra* (**A, B**), *Md-dsx* (**C, D**) and *Md-fru* (**E, F**) in abdomens (**A, C, E**) and heads (**B, D, F**) of flies with different RNAi treatments. Exons of *Md-tra*, *Md-dsx*, and *Md-fru* are presented along the X-axis. The names of the *Md-tra*, *Md-dsx*, and *Md-fru* exons follow published nomenclature (Hediger *et al*. 2004, 2010; Meier *et al*. 2013). Insets show female and male isoforms of *Md-tra*, *Md-dsx* and *Md-fru*, respectively. In *Md-tra*, Blue exons (E2b, E3) that contain premature stop codons are included in the male isoforms of *Md-tra* but excluded from the female isoforms. In *Md-fru*, red exons (s^f^ and f) that are contained in female isoforms have premature stop codons, but are excluded from the male isoforms. Exons (s,s^f^,f) upstream from an exon ‘c1’ of *Md-fru* are not included in the read depth coverage because they are not on the same scaffold in the genome assembly. To confirm that the *Md-tra*-RNAi treatment knocks down *Md-tra* expression, we examined the expression of *Md-tra* using RNA-seq coverage data collected from the abdomen and head of each of our four sample types (**A, B**). We expect the expression of *Md-tra* in females to be higher than in males because males produce a splice variant with a premature stop codon that is likely to be processed by the nonsense-mediated decay (NMD) pathway (Hediger *et al*. 2010; Kervestin and Jacobson 2012). In addition, the ovaries are expected to produce large amounts of *Md-tra* transcripts because *Md-tra* activity is necessary for maternal establishment of zygotic splicing of *Md-tra* via an auto-regulatory loop (Dübendorfer and Hediger 1998). In abdomen, normal females (GFP-RNAi treated genotypic females) do indeed express *Md-tra* approximately three times higher than normal males (genotypic males with either the GFP-RNAi or *Md-tra*-RNAi treatment). This high *Md-tra* expression in female abdomens might reflect the outcomes of strong ovarian expression. Importantly, *Md-tra* expression in sex-reversed males (genotypic females that are phenotypic males because of *Md-tra*-RNAi) was comparable to that of the genotypic males, not the normal females (**A**). This is likely because knock down of *Md-tra* by RNAi produces sex-reversed males that have functioning testes instead of ovaries. The *Md-tra* exons that are included in the functional, female-determining transcript were the highest expressed exons in phenotypic females (**A**), consistent with the production of the female-determining transcript in female ovaries (Hediger *et al*. 2010). We find that *Md-tra* is also differentially expressed between females and males in head, but the difference is much smaller than in abdomen (**B**). Notably, when we analyze the read mapping to *Md-tra* using DESeq2, expression is significantly higher in normal females than in genotypic (III^M^ #1) males. However, there is not a significant difference in *Md-tra* expression between sex-reversed males and either normal males or normal females. These results were observed after we excluded a sex-reversed male head sample that had an outlier expression profile (see Main Text). The lack of sexually dimorphic expression of *Md-tra* in head is consistent with minimal sex-biased expression in *Drosophila* and house fly heads (Goldman and Arbeitman 2007; Meisel *et al*. 2015). In addition, most somatic cells in *Drosophila* are sexually monomorphic as a result of cell autonomous sex determination in *Drosophila* (Robinett *et al*. 2010), suggesting that the same may be true for most cells in house fly heads. *Md-tra* regulates the splicing of at least two downstream genes, *Md-dsx* and *Md-fru*, which are both differentially spliced between females and males (Hediger *et al*. 2004, 2010; Meier *et al*. 2013). Only the female isoform of *Md-tra* is translated into a functional protein. In the presence of Md-Tra, *Md-dsx* is spliced into an isoform that promotes female morphological development. *Md-dsx* is spliced into an isoform that initiates male morphological development in the absence of Md-Tra (Hediger *et al*. 2004, 2010). *Md-fru* is spliced into a male-specific behavioral regulator in the absence of Md-Tra (Meier *et al*. 2013). The expression of *Md-dsx* and *Md-fru* in sex-reversed males was more similar to that of normal (genotypic) males (especially *Md-tra-*RNAi treated III^M^ males #2) than phenotypic females (**C-F**), confirming that *Md-tra* knock down affects the downstream genes in the sex determination pathway (Hediger *et al*. 2010; Meier *et al*. 2013). For example, *Md-dsx* expression in phenotypic males was higher than in phenotypic females, especially across male-specific exons (**C-D**), consistent with the expected effect of Md-TRA on *Md-dsx* splicing in females (Hediger *et al*. 2004, 2010). The expression of *Md-fru* was higher in head than in abdomen (**E-F**), consistent with its role as a behavioral regulator (Meier *et al*. 2013). Md-TRA regulates the splicing of *Md-fru* by promoting the production of splice variants with premature stop codons in females (Heinrichs *et al*. 1998; Meier *et al*. 2013). Sex-specific splicing of *Md-fru* occurs at the 5’ end of the transcript (Meier *et al*. 2013), but the 5’ end of *Md-fru* was not completely assembled and annotated in the reference genome. We therefore cannot test for differential splicing of *Md-fru* between males and females. However, we expect expression of *Md-fru* to be higher in males than females because the female splice variants will be removed by the NMD pathway. We indeed observe that *Md-fru* expression was much higher in the heads of GFP-RNAi treated genotypic males (III^M^ males #1; see Table 1) than GFP-RNAi treated normal females (**E-F**). However, in *Md-tra*-RNAi treated genotypic males (III^M^ males #2; see Table 1) and sex-reversed males, the expression of *Md-fru* is intermediate between females and GFP-RNAi treated genotypic males (**F**). A possible explanation is that RNAi knock down of *Md-tra* affects the expression or splicing of *Md-fru* in these flies (sex-reversed males and III^M^ males #2), but testing this hypothesis is beyond the scope of the work presented here.

**Supplementary Figure 8.**
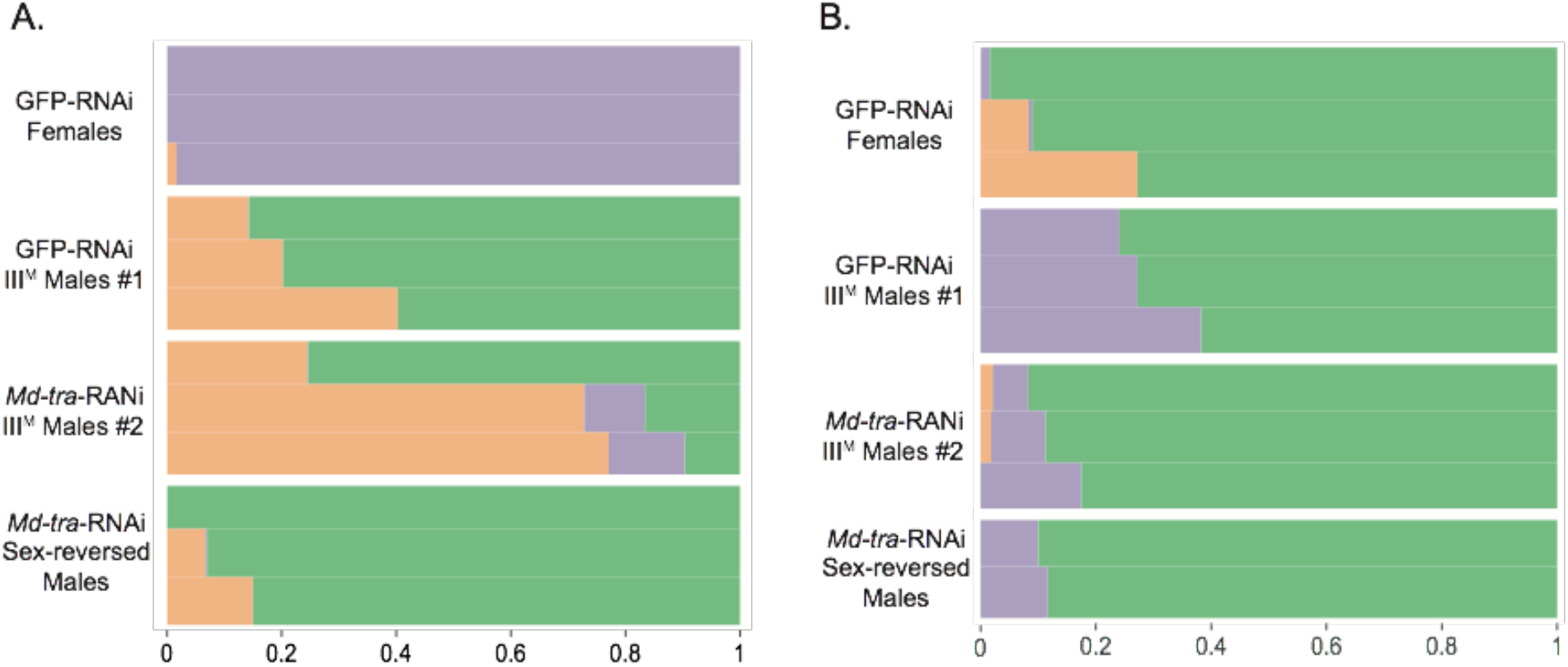
Grade of membership model (*K* = 3) for gene expression patterns of four types of dsRNA injected flies in abdomens (**A**) and heads (**B**). Each row is one replicate of each genotype-by-treatment combination. Each color represents the proportion of each replicate assigned to each of the three clusters.

**Supplementary Figure 9.**
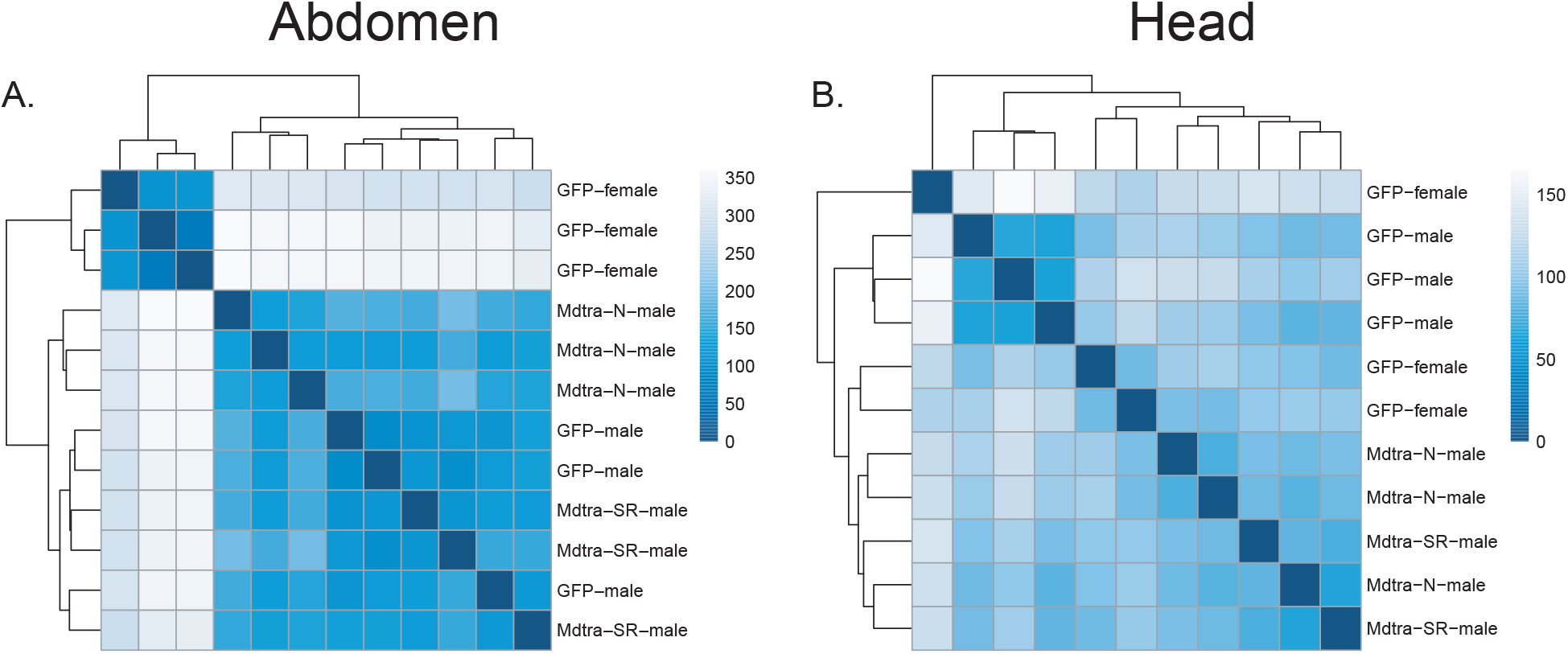
Hierarchical clustering of RNAi-treated flies in abdomen (**A**) and head (**B**). GFP-female stands for GFP-RNAi normal females, GFP-male for GFP-RNAi normal males #1, Mdtra-N-male for *Md-tra*-RNAi normal males #2, and Mdtra-SR-male for *Md-tra*-RNAi sex-reversed males.

**Supplementary Figure 10.**
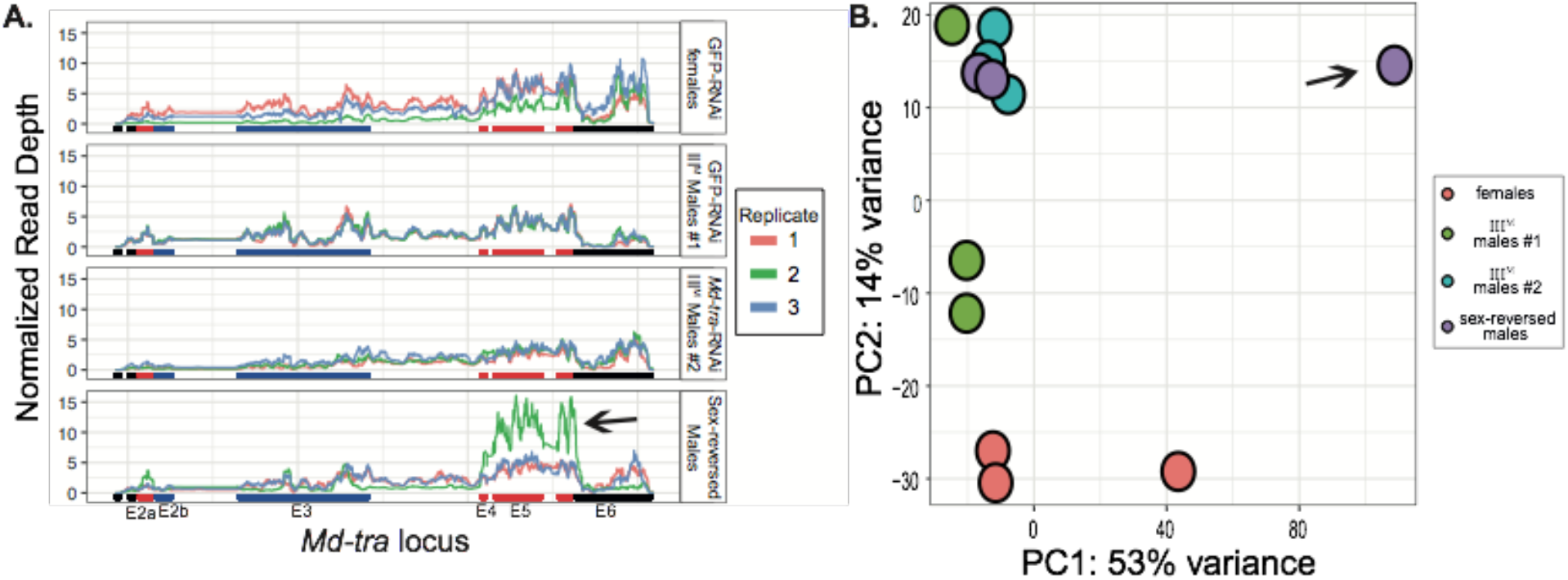
*Md-tra* expression (**A**) and PCA of global expression (**B**) of GPF-RNAi and *Md-tra*-RNAi individuals in heads. Arrows indicate the sex-reversed male head sample that we excluded from our analysis because of its outlier expression profile. Females are GFP-RNAi Normal Females; III^M^ males #1 are GFP-RNAi Normal Males; III^M^ males #2 are *Md-tra*-RNAi Normal Males; sex-reversed males are *Md-tra*-RNAi Sex-Reversed Males. SR stands for sex-reversed.

**Supplementary Figure 11.**
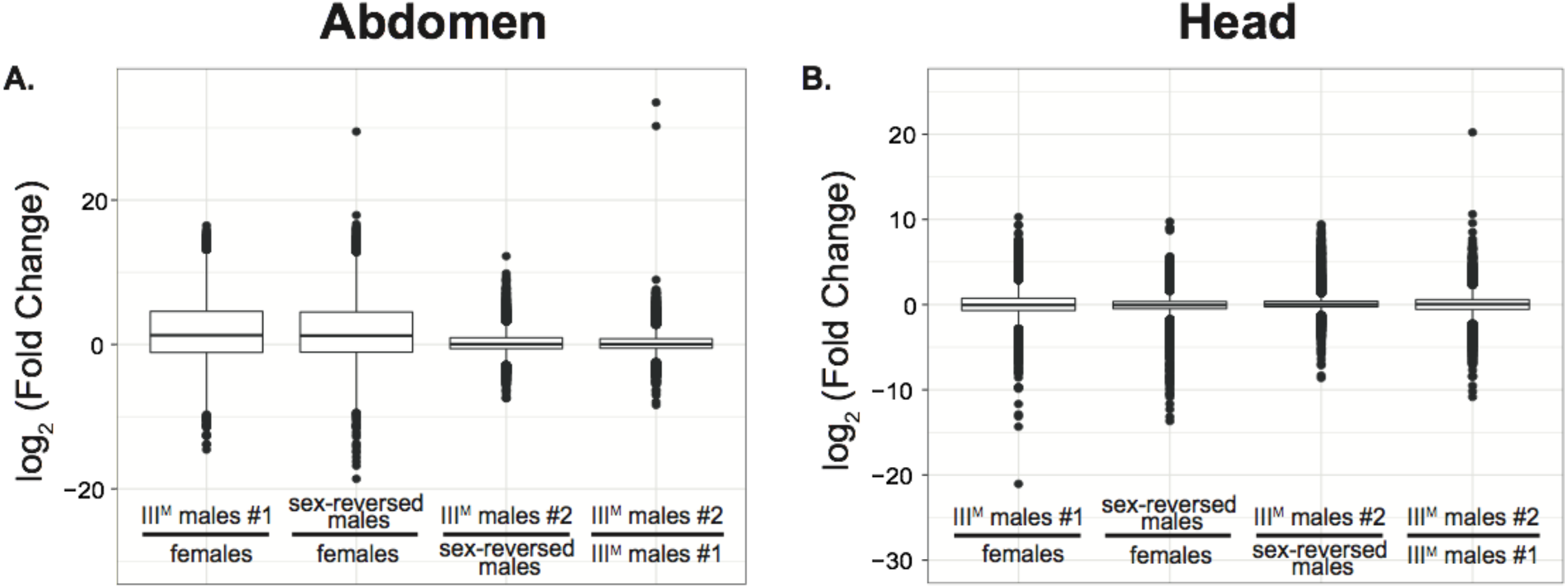
Boxplots show fold changes of gene expression among comparisons in abdomens (**A**) and heads (**B**). Outliers are included as points.

**Supplementary Figure 12.**
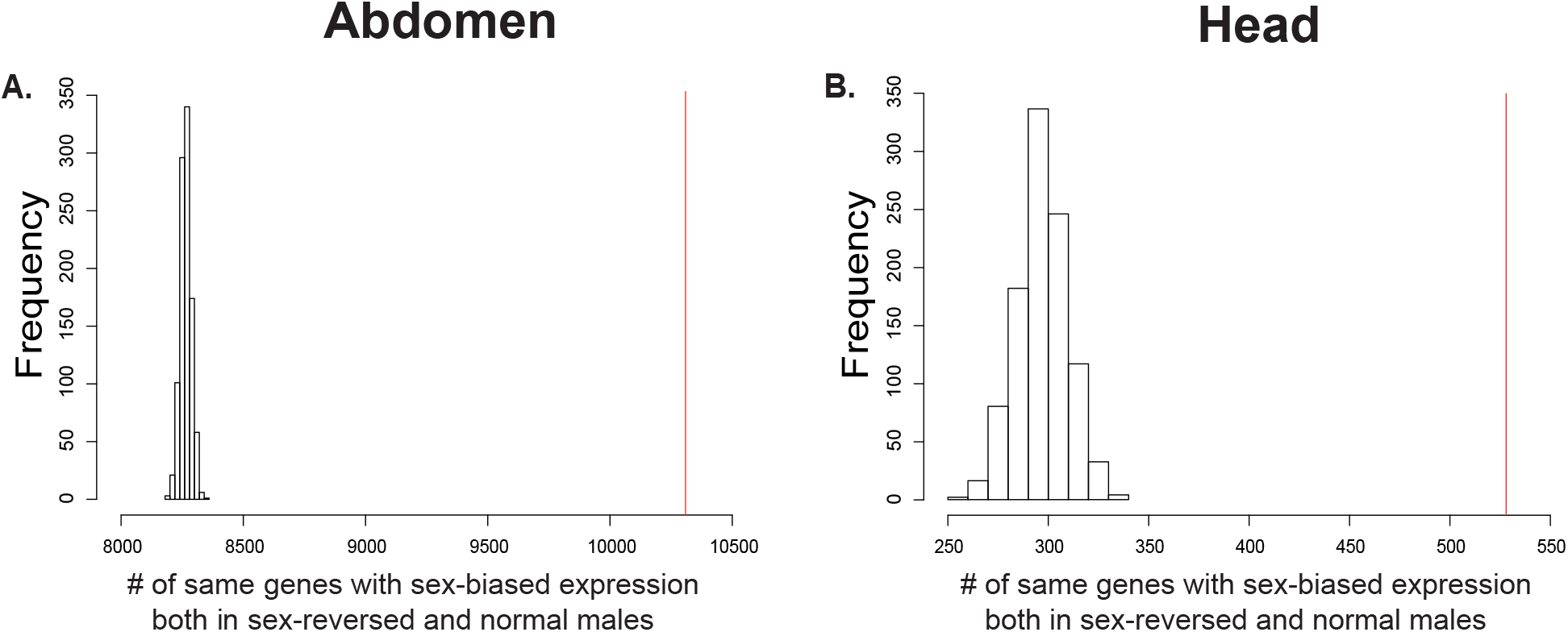
Permutation tests for whether the same genes have sex-biased expression both in sex-reversed males and normal males (III^M^ males #1) in abdomens (**A**) and heads (**B**). Histograms represent null distribution and red lines indicate the observed number of genes with the same sex-biased expression both in sex-reversed and normal males.

**Supplementary Table 1.** To confirm that the *Md-tra*-RNAi Number and percent of RNA-seq reads mapping to the house fly reference genome for females and each male with a different naturally occurring proto-Y chromosome.

| strain | sex | tissue | Reads mapped | Total reads | % mapped reads |
| --- | --- | --- | --- | --- | --- |
| bwbCS | male | abdomen | 28,797,051 | 53,216,164 | 54.11 |
| bwbCS | male | abdomen | 73,634,683 | 117,292,281 | 62.78 |
| bwbCS | male | abdomen | 34,387,641 | 46,794,091 | 73.49 |
| CS | male | abdomen | 23,436,748 | 43,892,080 | 53.40 |
| CS | male | abdomen | 47,225,783 | 73,027,117 | 64.67 |
| CS | male | abdomen | 29,656,467 | 39,569,021 | 74.95 |
| CSrab | male | abdomen | 26,931,208 | 44,702,325 | 60.25 |
| CSrab | male | abdomen | 10,639,480 | 17,154,576 | 62.02 |
| CSrab | male | abdomen | 39,105,284 | 51,590,908 | 75.80 |
| IsoCS | male | abdomen | 20,125,936 | 32,675,559 | 61.59 |
| IsoCS | male | abdomen | 5,422,571 | 8,475,350 | 63.98 |
| IsoCS | male | abdomen | 65,535,221 | 85,353,585 | 76.78 |
| IsoCS | female | abdomen | 49,248,555 | 61,591,761 | 79.96 |
| CSrab | female | abdomen | 24,268,638 | 33,301,977 | 72.87 |
| bwbCS | male | head | 41,466,785 | 55,597,781 | 74.58 |
| bwbCS | male | head | 36,041,171 | 50,840,587 | 70.89 |
| bwbCS | male | head | 62,840,805 | 80,615,028 | 77.95 |
| CS | male | head | 38,149,107 | 50,861,349 | 75.01 |
| CS | male | head | 19,919,990 | 27,227,824 | 73.16 |
| CS | male | head | 51,341,822 | 67,253,493 | 76.34 |
| CSrab | male | head | 56,191,834 | 75,651,390 | 74.28 |
| CSrab | male | head | 10,481,663 | 17,377,586 | 60.32 |
| CSrab | male | head | 55,120,600 | 73,965,574 | 74.52 |
| IsoCS | male | head | 17,332,681 | 27,276,701 | 63.54 |
| IsoCS | male | head | 28,841,223 | 39,570,265 | 72.89 |
| IsoCS | male | head | 62,213,672 | 82,095,892 | 75.78 |
| IsoCS | female | head | 49,089,361 | 63,708,772 | 77.05 |
| CSrab | female | head | 45,789,698 | 61,740,463 | 74.16 |
| CS | female | head | 45,947,520 | 57,865,112 | 79.40 |

**Supplementary Table 2.** Number and percent of RNA-seq reads mapping to the house fly reference genome for each sample from the RNAi experiment.

| Sample Name | Reads mapped | Total reads | % mapped reads |
| --- | --- | --- | --- |
| *Abdomen samples* |  |  |  |
| GFP-female-A2 | 91,396,275 | 106,578,289 | 85.76 |
| GFP-female-A3 | 79,790,013 | 93,362,685 | 85.46 |
| GFP-female-A4 | 67,507,547 | 78,662,688 | 85.82 |
| GFP-male-A1 | 30,825,624 | 41,147,995 | 74.91 |
| GFP-male-A2 | 43,102,152 | 57,473,417 | 74.99 |
| GFP-male-A3 | 43,544,801 | 59,178,052 | 73.58 |
| tra-N-A2 | 36,982,401 | 52,453,415 | 70.51 |
| tra-N-A3 | 68,463,509 | 96,288,121 | 71.10 |
| tra-N-A4 | 34,147,545 | 48,784,891 | 70.00 |
| tra-SR-A1 | 45,637,174 | 60,334,091 | 75.64 |
| tra-SR-A2 | 54,578,379 | 72,306,455 | 75.48 |
| tra-SR-A3 | 35,999,042 | 47,579,279 | 75.66 |
| *Head samples* |  |  |  |
| GFP-female-H2 | 41,898,942 | 53,096,665 | 78.91 |
| GFP-female-H3 | 54,190,305 | 69,827,731 | 77.61 |
| GFP-female-H4 | 68,366,760 | 90,002,468 | 75.96 |
| GFP-male-H1 | 86,270,347 | 106,498,553 | 81.01 |
| GFP-male-H2 | 76,486,314 | 96,639,044 | 79.15 |
| GFP-male-H3 | 105,491,479 | 132,492,819 | 79.62 |
| tra-N-H2 | 42,279,891 | 55,339,089 | 76.40 |
| tra-N-H3 | 22,602,895 | 29,743,740 | 75.99 |
| tra-N-H4 | 31,937,609 | 41,479,997 | 77.00 |
| tra-SR-H1 | 47,090,731 | 60,413,724 | 77.95 |
| tra-SR-H2 | 82,077,805 | 101,558,122 | 80.82 |
| tra-SR-H3 | 25,046,702 | 31,679,494 | 79.06 |

**Supplementary Table 3.** Differential expression between males with different genotypes. Counts of the number of genes that are expressed differentially (# Diff) and total genes expressed (#Genes) are shown, as well as the frequency of genes that are expressed differentially (Freq Diff). Y^M^ males are from the IsoCS strain; bwbCS Y^M^ males have a standard chromosome III from CS (bwbCS×CS males); III^M^ males are from either the CS or CSrab strain.

| **Tissue** | **Comparison** | **#Diff** | **#Genes** | **Freq Diff** |
| --- | --- | --- | --- | --- |
| **Abdomen** | Y<sup>M</sup> vs Y<sup>M</sup> with new chr III | 1159 | 11533 | 0.100 |
|  | Y<sup>M</sup> vs III<sup>M</sup> (CS) | 511 | 10344 | 0.049 |
|  | Y<sup>M</sup> vs III<sup>M</sup> (CSrab) | 479 | 9346 | 0.051 |
|  | III<sup>M</sup> (CS) vs III<sup>M</sup> (CSrab) | 196 | 10460 | 0.19 |
| **Head** | Y<sup>M</sup> vs Y<sup>M</sup> with new chr III | 878 | 11909 | 0.074 |
|  | Y<sup>M</sup> vs III<sup>M</sup> (CS) | 377 | 11845 | 0.032 |
|  | Y<sup>M</sup> vs III<sup>M</sup> (CSrab) | 525 | 12390 | 0.042 |
|  | III<sup>M</sup> (CS) vs III<sup>M</sup> (CSrab) | 739 | 12409 | 0.060 |

**Supplementary Table 4.** Differential expression between genotypic males and females with different RNAi treatments. Counts of the number of genes that are expressed differentially (#Diff) and total genes expressed (#Genes) are shown, as well as the frequency of genes that are expressed differentially (Freq Diff). Females are GFP-RNAi treated normal females; sex-reversed males are *Md-tra*-RNAi treated genotypic females; III^M^ males #1 are GFP-RNAi treated genotypic males; III^M^ males #2 are *Md-tra*-RNAi treated genotypic males.

| **Tissue** | **Comparison** | **#Diff** | **#Genes** | **Freq Diff** |
| --- | --- | --- | --- | --- |
| **Abdomen** | III<sup>M</sup> males #1 vs females | 11030 | 14993 | 0.736 |
|  | sex-reversed males vs females | 11005 | 14686 | 0.749 |
|  | III<sup>M</sup> males #2 vs sex-reversed males | 2867 | 13769 | 0.208 |
|  | III<sup>M</sup> males #2 vs III<sup>M</sup> males #1 | 2243 | 13162 | 0.170 |
| **Head** | III<sup>M</sup> males #1 vs females | 5077 | 13558 | 0.374 |
|  | sex-reversed males vs females | 735 | 12360 | 0.059 |
|  | III<sup>M</sup> males #2 vs sex-reversed males | 204 | 12959 | 0.016 |
|  | III<sup>M</sup> males #2 vs III<sup>M</sup> males #1 | 3260 | 13258 | 0.246 |

**Supplementary Table 5.** Genes with sex-biased expression in sex-reversed or genotypic males. Counts are the number of genes that belong to each column and row combination. Columns compare sex-reversed males and normal females. Rows compare genotypic (normal) males and normal females. Genes with male-biased (female-biased) expression are expressed at significantly different levels between the sexes. Genes with not female-biased (not male-biased) have log_2_M/F not greater (less) than zero.

| **Abdomen**<br>(# genes) |  | sex-reversed males vs females |  |  |  |
| --- | --- | --- | --- | --- | --- |
|  |  | male-biased | not female-biased | not male-biased | female-biased |
| genotypic male<br>(III<sup>M</sup> males #1)<br>vs<br>females | male-biased | 6136 | 407 | 19 | 1 |
|  | not female-biased | 483 | 1574 | 271 | 5 |
|  | not male-biased | 26 | 274 | 841 | 182 |
|  | female-biased | 5 | 18 | 277 | 4167 |
| **Head**<br>(# genes) |  | sex-reversed males vs females |  |  |  |
|  |  | male-biased | not female-biased | not male-biased | female-biased |
| genotypic male<br>(III<sup>M</sup> males #1)<br>vs<br>females | male-biased | 254 | 1897 | 380 | 4 |
|  | not female-biased | 50 | 2221 | 1177 | 34 |
|  | not male-biased | 13 | 1091 | 2660 | 110 |
|  | female-biased | 3 | 154 | 2045 | 267 |

**Supplementary Table 6.** Chromosomal distribution of discordant sex-biased genes in abdomen. The chromosomal distribution of discordant genes was compared to all genes in the genome. Genes that were not assigned to a chromosome were excluded. A Fisher’s exact test was performed to test for an excess of discordant sex-biased genes on each chromosome relative to genes on all other chromosomes.

|  |  | **Abdomen** |  |  |  |  |  |
| --- | --- | --- | --- | --- | --- | --- | --- |
|  |  | normal-up-discordant<br>(n-u-d) |  |  | sex-reversed-up-discordant<br>(sr-u-d) |  |  |
| **Chromosomes**<br>**(Muller**<br>**elements)** | **# genes**<br>**on chr** | **# genes** | **Odds**<br>**ratio** | **95% CI** | **# genes** | **Odds**<br>**ratio** | **95% CI** |
| **1(B)** | 2000 | 1 | 0.270 | 0.006 -<br>1.709 | 2 | 0.270 | 0.031 -<br>1.048 |
| **2(E)** | 2910 | 4 | 0.806 | 0.194 -<br>2.532 | 11 | 1.231 | 0.550 -<br>2.568 |
| **3(A)** | **2094** | **9** | **4.129** | **1.482 -**<br>**11.322** | **13** | **2.386** | **1.119 -**<br>**4.853** |
| **4(D)** | 2184 | 3 | 0.817 | 0.152 -<br>2.58 | 5 | 0.660 | 0.201 -<br>1.705 |
| **5(C)** | 2469 | 2 | 0.440 | 0.049 -<br>1.855 | 7 | 0.844 | 0.313 -<br>1.958 |
| **X(F)** | 45 | 0 | 0 | 0 -<br>57.539 | 0 | 0 | 0 -<br>27.459 |
| **Total** | 11702 | 19 |  |  | 38 |  |  |

**Supplementary Table 7.** Chromosomal distribution of discordant sex-biased genes in head. The chromosomal distribution of discordant genes was compared to all genes in the genome. Genes that were not assigned to a chromosome were excluded. A Fisher’s exact test was performed to test for an excess of discordant sex-biased genes on each chromosome relative to genes on the other chromosomes.

|  |  | Head |  |  |  |  |  |
| --- | --- | --- | --- | --- | --- | --- | --- |
|  |  | normal-up-discordant<br>(n-u-d) |  |  | sex-reversed-up-discordant<br>(sr-u-d) |  |  |
| Chromosomes<br>(Muller<br>elements) | # genes<br>on chr | # genes | Odds<br>ratio | 95% CI | # genes | Odds<br>ratio | 95% CI |
| **1(B)** | 1750 | 67 | 1.145 | 0.859 -<br>1.507 | 18 | 0.862 | 0.490 -<br>1.436 |
| **2(E)** | 2550 | 73 | 0.792 | 0.601 -<br>1.032 | 24 | 0.759 | 0.463 -<br>1.201 |
| **3(A)** | **1842** | **108** | **2.028** | **1.592 -**<br>**2.569** | **44** | **2.665** | **1.787 -**<br>**3.933** |
| **4(D)** | 1888 | 50 | 0.735 | 0.531 -<br>0.999 | 12 | 0.494 | 0.247 -<br>0.902 |
| **5(C)** | 2147 | 51 | 0.641 | 0.465 -<br>0.868 | 21 | 0.805 | 0.476 -<br>1.303 |
| **X(F)** | 34 | 1 | 0.858 | 0.021 -<br>5.144 | 0 | 0 | 0 -<br>9.947 |
| **Total** | 10211 | 350 |  |  | 119 |  |  |

